# Virus genomes from deep sea sediments expand the ocean megavirome and support independent origins of viral gigantism

**DOI:** 10.1101/469403

**Authors:** Disa Bäckström, Natalya Yutin, Steffen L. Jørgensen, Jennah Dharamshi, Felix Homa, Katarzyna Zaremba-Niedwiedzka, Anja Spang, Yuri I. Wolf, Eugene V. Koonin, Thijs J. G. Ettema

**Affiliations:** Department of Cell and Molecular Biology, Science for Life Laboratory, Uppsala University, Box 596, Uppsala SE-75123, Sweden; National Center for Biotechnology Information, National Library of Medicine. National Institutes of Health, Bethesda, MD 20894, USA; Department of Biology, Centre for Geobiology, University of Bergen, N-5020 Bergen, Norway; NIOZ, Royal Netherlands Institute for Sea Research, Department of Marine Microbiology and Biogeochemistry, and Utrecht University, P.O. Box 59, NL-1790 AB Den Burg, The Netherlands

**Author notes:** These authors have contributed equally (Disa Bäckström, Natalya Yutin).

## Abstract

The Nucleocytoplasmic Large DNA Viruses (NCLDV) of eukaryotes (proposed order ”Megavirales”) include the families *Poxviridae, Asfarviridae, Iridoviridae, Ascoviridae, Phycodnaviridae, Marseilleviridae,* and *Mimiviridae*, as well as still unclassified Pithoviruses, Pandoraviruses, Molliviruses and Faustoviruses. Several of these virus groups include giant viruses, with genome and particle sizes exceeding those of many bacterial and archaeal cells. We explored the diversity of the NCLDV in deep-sea sediments from the Loki’s Castle hydrothermal vent area. Using metagenomics, we reconstructed 23 high quality genomic bins of novel NCLDV, 15 of which are closest related to Pithoviruses, 5 to Marseilleviruses, 1 to Iridoviruses, and 2 to Klosneuviruses. Some of the identified Pitho-like and Marseille-like genomes belong to deep branches in the phylogenetic tree of core NCLDV genes, substantially expanding the diversity and phylogenetic depth of the respective groups. The discovered viruses have a broad range of apparent genome sizes including putative giant members of the family *Marseilleviridae*, in agreement with multiple, independent origins of gigantism in different branches of the NCLDV. Phylogenomic analysis reaffirms the monophyly of the Pitho-Irido-Marseille branch of NCLDV. Similarly to other giant viruses, the Pitho-like viruses from Loki’s Castle encode translation systems components. Phylogenetic analysis of these genes indicates a greater bacterial contribution than detected previously. Genome comparison suggests extensive gene exchange between members of the Pitho-like viruses and *Mimiviridae*. Further exploration of the genomic diversity of “Megavirales” in additional sediment samples is expected to yield new insights into the evolution of giant viruses and the composition of the ocean megavirome.

**Importance:** Genomics and evolution of giant viruses is one of the most vigorously developing areas of virus research. Lately, metagenomics has become the main source of new virus genomes. Here we describe a metagenomic analysis of the genomes of large and giant viruses from deep sea sediments. The assembled new virus genomes substantially expand the known diveristy of the Nucleo-Cytoplasmic Large DNA Viruses of eukaryotes. The results support the concept of independent evolution of giant viruses from smaller ancestors in different virus branches.

## Introduction

The nucleocytoplasmic large DNA viruses (NCLDV) comprise an expansive group of viruses that infect diverse eukaryotes (1). Most of the NCLDV share the defining biological feature of reproducing (primarily) in the cytoplasm of the infected cells as well as several genes encoding proteins involved in the key roles in virus morphogenesis and replication, leading to the conclusion that the NCLDV are monophyletic, that is, evolved from a single ancestral virus (2, 3). As originally defined in 2001, the NCLDV included 5 families of viruses: *Poxviridae*, *Asfarviridae, Iridoviridae*, *Ascoviridae*, and *Phycodnaviridae* (2). Subsequent isolation of viruses from protists has resulted in the stunning discovery of giant viruses, with genome sizes exceeding those of many bacteria and archaea (4-8). The originally discovered group of giant viruses has formed the family *Mimiviridae* (9-13). Subsequently, 3 additional other groups of giant viruses have been identified, namely, Pandoraviruses (14-16);Pithoviruses, Cedratviruses and Orpheovirus (hereafter, the latter 3 groups of related viruses are collectively referred to as the putative family ”Pithoviridae”) (17-19), and *Mollivirus sibericum* (20), along with two new groups of NCLDV with moderate-sized genomes, the family *Marseilleviridae* (21, 22), and Faustoviruses (23, 24). Most of the NCLDV have icosahedral virions composed of a double jelly roll major capsid proteins but Poxviruses have distinct brick-shaped virions, ascoviruses have ovoid virions, Mollivirus has a spherical virion, finally, Pandoraviruses and Pithoviruses have unusual, amphora-shaped virions.. The Pithovirus virions are the largest among the currently known viruses. Several of the recently discovered groups of NCLDV are likely to eventually become new families in particular, the putative ; family ”Pithoviridae” (25), and reclassification of the NCLDV into a new virus order “Megavirales” has been proposed (26, 27).

Phylogenomic reconstruction of gene gain and loss events resulted in mapping about 50 genes that are responsible for the key viral functions to the putative last common ancestor of the NCLDV, reinforcing the conclusion on their monophyly (3, 28). However, detailed phylogenetic analysis of these core genes of the NCLDV has revealed considerable evolutionary complexity including numerous cases of displacement of ancestral genes with homologs from other sources, and even some cases of independent capture of homologous genes (29). The genomes of the NCLDV encompass from about 100 (some iridoviruses) to nearly 2500 genes (pandoraviruses) that, in addition to the 50 or so core genes, include numerous genes involved in various aspects of virus-host interaction, in particular, suppression of the host defense mechanisms, as well as many genes for which no function could be identified (1, 30).

The NCLDV include some viruses that are agents of devastating human and animal diseases, such as smallpox virus or African swine fever virus (31, 32), as well as viruses that infect algae and other planktonic protists and are important ecological agents (12, 33-35). Additionally, NCLDV elicit strong interest of many researchers due to their large genome size which, in the case of the giant viruses, falls within the range of typical genome size of bacteria and archaea. This apparent exceptional position of the giant viruses in the virosphere, together with the fact that they encode multiple proteins that are universal among cellular organisms, in particular, translation system components, has led to provocative scenarios of the origin and evolution of giant viruses. It has been proposed that the giant viruses were descendants of a hypothetical, probably, extinct fourth domain of cellular life that evolved via drastic genome reduction, and support of this scenario has been claimed from phylogenetic analysis of aminoacyl-tRNA synthetases encoded by giant viruses (5, 26, 36-40). However, even apart from the conceptual difficulties inherent in the postulated cell to virus transition (41, 42), phylogenetic analysis of expanded sets of translation-related proteins encoded by giant viruses has resulted in tree topologies that were poorly compatible with the fourth domain hypothesis but rather suggest piecemeal acquisition of these genes, likely, from different eukaryotic hosts (43-46).

More generally, probabilistic reconstruction of gene gains and losses during the evolution of the NCLDV has revealed a highly dynamic evolutionary regime (3, 28, 29, 45, 46) that has been conceptualized in the so-called genomic accordion model under which virus evolution proceeds via alternating phases of extensive gene capture and gene loss (47, 48). In particular, in the course of the NCLDV evolution, giant viruses appear to have evolved from smaller ones on multiple, independent occasions (45, 49, 50).

In recent years, metagenomics has become the principal route of new virus discovery (51-53). However, in the case of giant viruses, *Acanthamoeba* co-culturing has remained the main source of new virus identification, and this methodology has been refined to allow for high-throughput giant virus isolation (54, 55). To date, over 150 species of giant viruses have been isolated from various environments, including water towers, soil, sewage, rivers, fountains, seawater, and marine sediments (56). The true diversity of giant viruses is difficult to assess, but the explosion of giant virus discovery during the last ten years, and large scale metagenomic screens of viral diversity indicates that a major part of the Earth’s virome remains unexplored (57). The core genes of the NCLDV can serve as baits for screening environmental sequences, and pipelines have been developed for large scale screening of metagenomes (56, 58). Although these efforts have given indications of the presence of uncharacterized giant viruses in samples from various environments, few of these putative novel viruses can be characterized due to the lack of genomic information. Furthermore, giant viruses tend to be overlooked in viral metagenomic studies since samples are typically filtered according to the preconception of typical virion sizes (52).

To gain further insight into the ecology, evolution, and genomic content of giant viruses, it is necessary to retrieve more genomes, not simply establish their presence by detection of single marker genes. Metagenomic binning is the process of clustering environmental sequences that belong to the same genome, based on features such as base composition and coverage. Binning has previously been used to reconstruct the genomes of large groups of uncharacterized bacteria and archaea in a culture-independent approach (59, 60). Only one case of binning has been reported for NCLDV, when the genomes of the Klosneuviruses, distant relatives of the Mimiviruses, were reconstructed from a simple wastewater sludge metagenome (46). More complex metagenomes from all types of environments remain to be explored. However, standard methods for screening and binning of NCLDV have not yet been developed, and sequences of these viruses can be difficult to classify because of substantial horizontal gene transfer from bacteria and eukaryotes (13, 29, 43, 49), and also because a large proportion of the NCLDV genes (known as ORFans) have no detectable homologs (25, 30).

We identified NCLDV sequences in deep sea sediment metagenomes from Loki’s Castle, a sample site that has been previously shown to be rich in uncharacterized prokaryotes (61, 62) (Dharamshi et. al. 2018 (submitted)). The complexity of the data and genomes required a combination of different binning methods, assembly improvement by reads profiling, and manual refinement of each bin to minimize contamination with non-viral sequences. As a result, 23 high quality genomic bins of novel NCLDV were reconstructed, including, mostly, distant relatives of ”Pithoviridae”, Orpheovirus, and *Marseilleviridae*, as well as two relatives of Klosneuviruses. These findings substantially expand the diversity of the NCLDV, in particular, the Pitho-Irido-Marseille (PIM) branch, further support the scenario of independent evolution of giant viruses from smaller ones in different branches of the NCLDV, and provide an initial characterization of the ocean megavirome.

## Materials and Methods

### Sampling and metagenomic sequencing

In the previous studies of microbial diversity in the deep sea sediments, samples were retrieved from three sites about 15 km north east of the Loki's castle hydrothermal vent field (Table S1 of Additional File 1), by gravity (GS10_GC14, GS08_GC12) and piston coring (GS10_PC15) (61, 63, 64).

DNA was extracted and sequenced, and metagenomes were assembled as part of the previous studies ((61) for GS10_GC14, Dharamshi et. al. 2018 (submitted) for GS08_GC12 and GS10_PC15), resulting in the assemblies LKC75, KR126, K940, K1000, and K1060. Contiguous sequences (contigs) <u>longer than</u> 1kb were selected for further processing.

### Identification of viral metagenomic sequences

Protein sequences of the metagenomic contigs were predicted using Prodigal v.2.6.3 (65), in the metagenomics mode. A collection of DNA polymerase family B (DNAP) sequences from 11 NCLDV was used to query the metagenomic protein sequence with BLASTP ((66), Table S1 of Additional File 1). The BLASTP hits were filtered according to e-value (maximum 1e^-5^), alignment length (at least 50% of the query length) and identity (greater than 30%). The sequences were aligned using MAFFT-LINSI (67). Reference NCLDV DNAP sequences were extracted from the NCVOG collection (28). Highly divergent sequences and those containing large gaps inserts were removed from the alignment, followed by re-alignment. The terminal regions of the alignments were trimmed manually using Jalview (68), and internal gaps were removed using trimAl (v.1.4.rev15, (69)) with the option “gappyout”. IQTree version 1.5.0a (70) was used to construct maximum likelihood phylogenies with 1000 ultrafast bootstrap replications (71). The built-in model test (72) was used to select the best evolutionary model according to the Bayesian information criterion (LG+F+I+G4; Figure S1 of Additional File 1). Contigs belonging to novel NCLDVs were identified and used for binning.

### Composition-based binning (ESOM)

All sequences of the assemblies KR126, K940, K1000 and K1060 were split into fragments of minimum 5 or 10 kb length at intervals of 5 or 10 kb, and clustered by tetranucleotide frequencies using Emergent Self Organizing maps (ESOM, (73)), generating one map per assembly. Bins were identified by viewing the maps using Databionic ESOM viewer (http://databionic-esom.sourceforge.net/), and manually choosing the contigs clustering together with the putative NCLDV contigs in an “island” (Figure S3 of Additional File 1).

### Differential coverage binning of metagenomic contigs

Differential coverage (DC) bins were generated for the KR126, K940, K1000, and K1060 metagenomes, according to Dharamshi et. al. 2018 (submitted). Briefly, Kallisto version 0.42.5 (74) was used to get the differential coverage data of each read mapped onto each focal metagenome, that was used by CONCOCT version 0.4.1 to collect sequences into bins (75). CONCOCT was run with three different contig size thresholds: 2kb, 3kb, and 5kb, and longer contigs were cut up into smaller fragments (10 kb), to decrease coverage and compositional bias, and merged again after CONCOCT binning (See Dharamshi et. al. 2018 (submitted) for further details). Bins containing contigs with the viral DNAP were selected and refined in mmgenome (76)). Finally, to resolve overlapping sequences in the DC bins, the reads of each bin were extracted using seqtk (version 1.0-r82-dirty, https://github.com/lh3/seqtk) and the reads mapping files generated for mmgenome, and reassembled using SPAdes (3.6.0, multi-cell, --careful mode, (77)). Bins from KR126 had too low coverage and quality, and were discarded from further analysis.

### Co-assembly binning of metagenomic contigs

CLARK (78), a program for classification of reads using discriminative k-mers, was used to identify reads belonging to NCLDV in the metagenomes. A target set of 10 reference genomes that represented Klosneuviruses, *Marseilleviridae*, and ”Pithoviridae” (Table S2 of Additional File 1), as well as the 29 original bins, were used to make a database of spaced k-mers which CLARK used to classify the reads of the K940, K1000 and K1060 metagenomes (full mode, k-mer size 31). Reads classified as related to any of the targets were extracted and the reads from all three metagenomes were pooled and reassembled using SPAdes (3.9.0, (77)). Because CLARK removes not-discriminatory k-mers, the reads for sequences that are similar between the bins might not have been included. Therefore, the reads from each original bin that were used for the first set reassemblies, were also included, and pooled with the CLARK-classified reads before reassembly.

Four SPAdes modes were tested: metagenomic (--meta), sincle-cell (--sc), multi-cell (default), and multi-cell careful (--careful). The quality of the assemblies was tested by identifying the contigs containing NCVOG0038 (DNA polymerase), using BLASTP (66). The multi-cell careful assembly had the longest DNAP-containing contigs and was used for CONCOCT binning.

CONCOCT was run as above, only using reads from the co-assembly as input. Bins containing NCVOG0038 were identified by BLASTP. The smaller the contig size threshold, the more ambiguous and potentially contaminating sequences were observed, so the CONCOCT 5 kb run was chosen to extract and refine new bins. The bins were refined by using mmgenome as described below.

### Quality assessment and refinement of metagenomic NCLDV bins

General sequence statistics were calculated by Quast (v. 3.2, (79)). Barrnap (v 0.8; (80)) was used to check for the presence of rRNA genes, with a length threshold of 0.1. Prokka (v1.12, (79)) was used to annotate open reading frames (ORFs) of the raw bins. Megavirus marker gene presence in each metagenomic bin was estimated by using the micomplete pipeline (https://bitbucket.org/evolegiolab/micomplete) and a set of the 10 conserved NCLDV genes (Table S3 of Additional File 1). This information was used to assess completeness and redundancy. Presence of more than one copy of each marker gene was considered an indication of potential contamination or the presence of more than one viral genome per bin, and such bins were further refined.

Mmgenome was used to manually refine the metagenomic bins by plotting coverage and GC-content, showing reads linkage, and highlighting contigs with marker genes (76). Overlap between the ESOM binned contigs and the DC bins was also visualized. Bins containing only one genome were refined by removing contigs with different composition and coverage. In cases when several genomes were represented in the same CONCOCT bin, they were separated into different bins when distinct clusters were clearly visible (see the Supplementary Materials of Additional File 1 for examples of the refining process). Reads linkage was determined by mapping the metagenomic reads onto the assembly using bowtie2 (version 2.3.2, (81)), samtools (version 1.2, (82)) to index and convert the mapping file into bam format, and finally a script provided by the CONCOCT suite to count the number of read pairs that were mapping to the first or last 1 kb of two different contigs (bam_to_linkage.py, --regionlength 1000).

Diamond aligner Blastp (83) was used to query the protein sequences of the refined bins against the NCBI non-redundant protein database (latest date of search: Febuary 13 2018), with maximum e-value 1e^-5^. Taxonomic information from the top BLASTP hit for each gene was used for taxonomic filtering. Contigs were identified as likely contaminants and removed if they had 50% or more bacterial or archaeal hits compared to no significant hits, and no viral or eukaryotic hits.

The assemblies of the DC and CA bins were compared by aligning the contigs with nucmer (part of MUMmer3.23,(84)) and an in-house script for visualization (see Additional File 1 for more details).

### Assessment of NCLDV diversity

Environmental sequences, downloaded in March 2017 from TARA Oceans ((85), https://www.ebi.ac.uk/ena/about/tara-oceans-assemblies), and EarthVirome ((57), available at https://img.jgi.doe.gov/vr/) were combined with the metagenomic sequences from Loki’s Castle (Table S1 of Additional file 1) and screened for sequences related to the Loki’s Castle NCLDVs using BLASTP search with the bin DNAP sequences as queries. The BLASTP hits were filtered according to e-value (maximum 1e^-5^), HSP length (at least 50% of the query length) and identity above 30%. The sequences were extracted using blastdbcmd, followed by alignment and phylogenetic tree reconstruction as described above (Figure 1).

**Figure 1.**
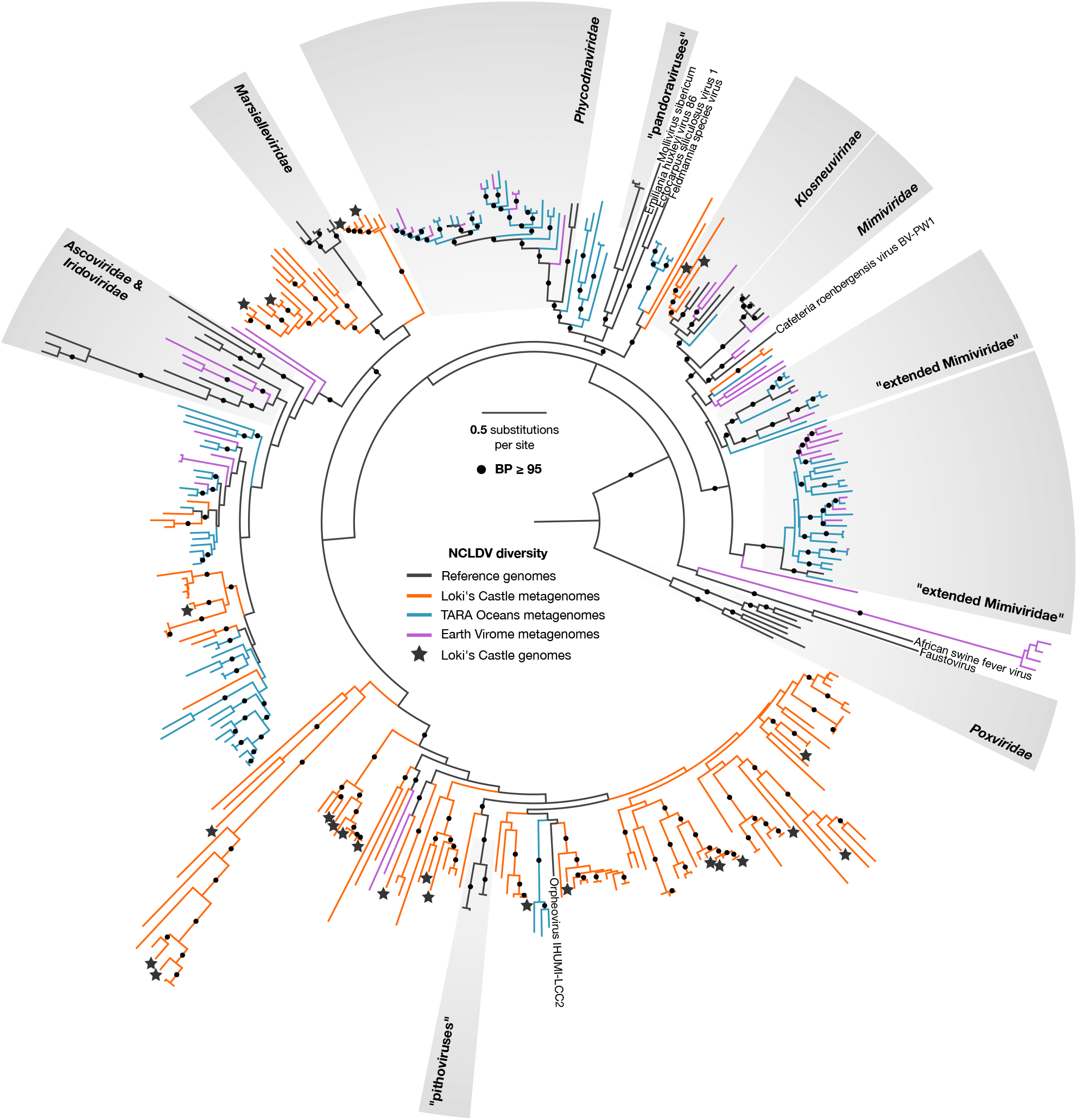
Diversity of the NCLDV DNAP sequences in the Loki’s Castle sediment metagenomes (orange), and the TARA oceans (turquoise), and EarthVirome (purple) databases. Reference sequences are shown in black. The binned NCLDV genomes are marked with a star. Branches with bootstrap values above 95 are marked with a black circle. The maximum likelihood phylogeny was constructed as described under Methods.

### Sequence annotation and phylogenetic analysis

The sequences of the selected bins were translated with MetaGeneMark (86). tRNA genes were predicted using tRNAscan-SE online (87). Predicted proteins were annotated using their best hits to NCVOG, cdd, and *nr* databases. In addition, Pitho-, Marseille-, Iridovirus-related bins were annotated using protein clusters constructed as described below. Reference sequences were collected from corresponding NCVOG and cdd profiles, and from GenBank, using BLASTP searches initiated from the Loki’s Castle NCLDV proteins. Reference sequences for Loki’s Castle virophages were retrieved by BLAST and tBLASTn searches against genomic (nr) and metagenomic (environmental wgs) parts of GenBank, with the predicted Loki’s Castle virophage MCP as queries. The retrieved environmental virophage genome fragments were translated with MetaGeneMark. Homologous sequences were aligned using MUSCLE (88). For phylogenetic reconstruction, gapped columns (more than 30% of gaps) and columns with low information content were removed from the alignments (89); the filtered alignments were used for tree reconstructions using FastTree (90). The alignments of three conserved NCLDV proteins were concatenated and used for phylogenetic analysis with PhyML ((91), http://www.atgc-montpellier.fr/phyml-sms/) The best model identified by PhyML was LG +G+I+F (LG substitution model, gamma distributed site rates with gamma shape parameter estimated from the alignment; fraction of invariable sites estimated from the alignment; and empirical equilibrium frequencies).

### Protein sequence clusters

Two sets of viral proteins, Pitho-Irido-Marseillevirus group (PIM clusters, ftp://ftp.ncbi.nih.gov/pub/yutinn/Loki_Castle_NCLDV_2018/PIM_clusters/) and NCLDV (NCLDV clusters, ftp://ftp.ncbi.nih.gov/pub/yutinn/Loki_Castle_NCLDV_2018/NCLDV_clusters/) were used separately to obtain two sets of protein clusters, using an iterative clustering and alignment procedure, organized as follows

- ***initial sequence clustering:*** Initially, sequences were clustered using UCLUST (92) with the similarity threshold of 0.5; clustered sequences were aligned using MUSCLE, singletons were converted to pseudo-alignments consisting of just one sequence. Sites containing more than 67% of gaps were temporarily removed from alignments and the pairwise similarity scores were obtained for clusters using HHSEARCH. Scores for a pair of clusters were converted to distances [the *d_A,B_* = -log(*s_A,B_*/min(*s_A,A_*,*s_B,B_*)) formula was used to convert scores ***s*** to distances ***d***)] a UPGMA guide tree was produced from a pairwise distance matrix. A progressive pairwise alignment of the clusters at the tree leaves was constructed using HHALIGN (93), resulting in larger clusters. The procedure was repeated iteratively, until all sequences with detectable similarity over at least 50% of their lengths were clustered and aligned together. Starting from this set of clusters, several rounds of the following procedures were performed.
- ***cluster merging and splitting:*** PSI-BLAST (94) search using the cluster alignments to construct Position-Specific Scoring Matrices (PSSMs) was run against the database of cluster consensus sequences. Scores for pairs of clusters were converged to a distance matrix as described above; UPGMA trees were cut using at the threshold depth; unaligned sequences from the clusters were collected and aligned together. An approximate ML phylogenetic tree was constructed from each of these alignments using FastTree (WAG evolutionary model, gamma-distributed site rates). The tree was split into subtrees so as to minimize paralogy and maximize species (genome) coverage. Formally, for a subtree containing *k* genes belonging to *m* genomes (*k* ≥ *m*) in the tree with the total of *n* genomes (*n* ≥ *m*) genomes, the “autonomy” value was calculated as (*m*/*k*)(*m*/*n*)(*a*/*b*)^1/6^ (where *a* is the length of the basal branch of the subtree and *b* is the length of the longest internal branch in the entire tree). This approach gives advantage to subtrees with the maximum representation of genomes, minimum number of paralogs and separated by a long internal branch. If a subtree with the maximum autonomy value was different from the complete tree, it was pruned from the tree, recorded as a separate cluster, and the remaining tree was analyzed again.
- ***cluster cutting and joining:*** Results of PSI-BLAST search whereby the cluster alignments were used as PSSMS and run against the database of cluster consensus sequences were analyzed for instances where a shorter cluster alignment had a full-length match to a longer cluster containing fewer sequences. This situation triggered cutting the longer alignment into fragments matching the shorter alignment(s). Alignment fragments were then passed through the merge-and-split procedure described above. If the fragments of the cluster that was cut did not merge into other clusters, the cut was rolled back, and the fragments were joined.
- ***cluster mapping and realigning:*** PSI-BLAST search using the cluster alignments as PSSMswas run against the original database. Footprints of cluster hits were collected, assigned to their respective highest-scoring query cluster and aligned, forming the new set of clusters mirroring the original set.
- ***post-processing:*** The PIM group clusters were manually curated and annotated using the NCVOG, CDD and HHPRED matches as guides. For the NCLDV clusters, the final round clusters with strong reciprocal PSI-BLAST hits and with compatible phyletic patters (using the same autonomy value criteria as described above) were combined into clusters of homologs that maximized genome representation and minimized paralogy. The correspondence between the previous version of NCVOGs and the current clusters was established by running PSI-BLAST with the NCVOG alignments as PSSMs against the database of cluster consensus sequences.

### Genome similarity dendrogram

Binary phyletic patterns of the NCLDV clusters (whereby 1 indicates a presence of the given cluster in the given genome) were converted to intergenomic distances as follows: *d_X,Y_* = -log(*N_X,Y_*/(*N_X_N_Y_*)^1/2^) where *N_X_* and *N_Y_* are the number of COGs present in genomes *X* and *Y* respectively and *N_X,Y_* is the number of COGs shared by these two genomes. A genome similarity dendrogram was reconstructed from the matrix of pairwise distances using the Neighbor-Joining method (95).

### Conserved motif search

The sequences from the LCV genomic bins were searched for potential promoters as follows. For every predicted ORF, ‘upstream’ genome fragments (from 250 nucleotides upstream to 30 nucleotides downstream of the predicted translation start codons) were extracted; short fragments (less than 50 nucleotides) were excluded; the resulting sequence sets were searched for recurring ungapped motifs using MEME software, with motif width set to either 25, 12, or 8 nucleotides (96). The putative LCV virophage promoter was used as a template to search upstream fragments of LCMiAC01 and LCMiAC02 with FIMO online tool (96). The motifs were visualized using the Weblogo tool (97).

### Data availability

The nucleotide sequences reported in this work have been deposited in GenBank under the accession numbers X00001-X0000N.

## Results

### Putative NCLDV in the Loki’s Castle metagenome

Screening of the Loki’s Castle metagenomes, for NCLDV DNA polymerase sequences revealed remarkable diversity (Figure 1, Figure S2, Additional File 1). Using two main binning approaches, namely, differential coverage binning (DC), and co-assembly binning (CA) (Figure 2), we retrieved 23 high quality bins of putative new NCLDVs (Table 1). The highest quality bins were identified by comparing the DC and the CA bins, based on decreasing the total number of contigs and the number of contigs without NCLDV hits, while preserving completeness (Additional File 1, Table S6).

**Figure 2.**
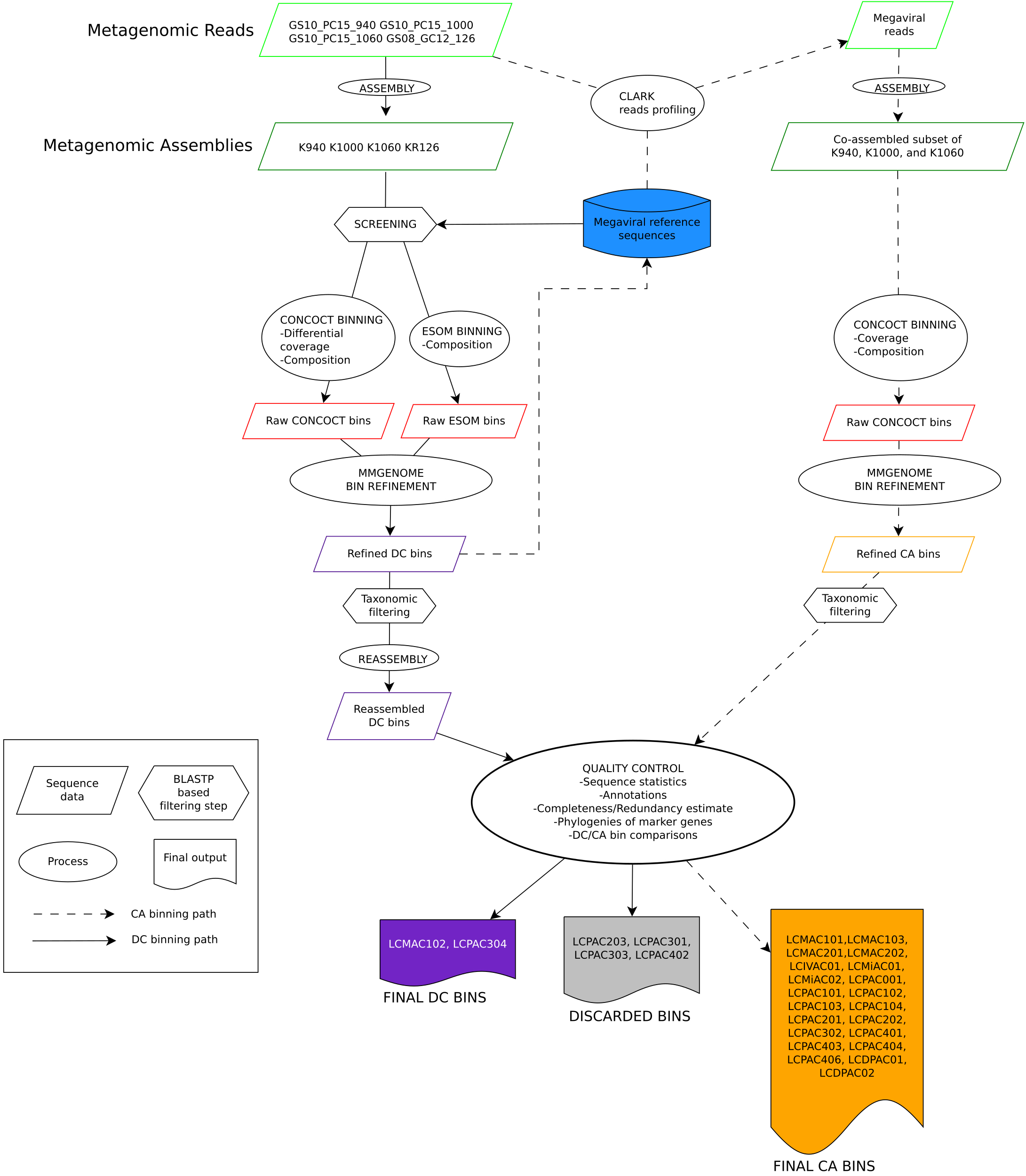
Flowchart of the metagenomic binning procedures. Two main binning approaches were used: differential coverage binning (DC), and co-assembly binning (CA). DC: Reads from four different samples were assembled into four metagenomes. The metagenomes were screened for NCLDV DNAP, and contigs were binned with CONCOCT and ESOM. The raw CONCOCT and ESOM bins were combined and refined using Mmgenome. The refined bins were put through taxonomic filtering, keeping only the contigs encoding at least one NCLDV gene, and finally, reassembled. CA: A database containing the refined DC bins and NCLDV reference genomes was used to create profiles to extract reads from the metagenomes. The reads were combined and co-assembled. This step was followed by CONCOCT binning, Mmgenome bin refinement and taxonomic filtering. Finally, the DC bins and CA bins were annotated and the best bins were chosen by comparing sequence statistics, completeness and redundancy of marker genes, and marker gene phylogenies (see Additional File 1 for details).

**Table 1.** The 23 NCLDV bins from Loki’s Castle.

| bin/virus |  |  | # of<br>contigs | min<br>contig<br>length, nt | max<br>contig<br>length, nt | total contig<br>length, nt | # of<br>predicted<br>proteins | MCP<sup>a</sup> | DNAp | ATP | RNAP<sup>A</sup> | RNAP<sup>B</sup> | D5hel | A18hel | VLTF3 | VLTF2 | RNAP5 | Erv1 | RNAlig | TopoII | FLAP | TFIIB |
| --- | --- | --- | --- | --- | --- | --- | --- | --- | --- | --- | --- | --- | --- | --- | --- | --- | --- | --- | --- | --- | --- | --- |
| LCPAC001 | Pitho-like |  | 12 | 8088 | 60499 | 249064 | 227 | 1 | 1 | 0 | 1 | 1 | 0 | 1 | 0 | 0 | 0 | 2 | 1 | 1 | 0 | 0 |
| LCPAC101 | Pitho-like |  | 26 | 6043 | 46492 | 466072 | 373 | 1 | 1 | 0 | 1 | 1 | 1 | 1 | 0 | 1 | 1 | 0 | 1 | 1 | 1 | 1 |
| LCPAC102 | Pitho-like |  | 12 | 6510 | 44810 | 285593 | 229 | 1 | 1 | 0 | 1 | 1 | 0 | 0 | 0 | 1 | 0 | 3 | 1 | 0 | 1 | 1 |
| LCPAC103 | Pitho-like |  | 17 | 5380 | 23680 | 204602 | 186 | 1 | 1 | 0 | 1 | 1 | 1 | 1 | 1 | 1 | 1 | 0 | 1 | 1 | 0 | 1 |
| LCPAC104 | Pitho-like |  | 4 | 6208 | 129049 | 218903 | 194 | 1 | 1 | 0 | 1 | 1 | 1 | 1 | 1 | 1 | 1 | 1 | 1 | 1 | 1 | 1 |
| LCPAC201 | Pitho-like |  | 11 | 5186 | 168698 | 428611 | 327 | 1 | 1 | 0 | 1 | 2 | 1 | 1 | 1 | 1 | 1 | 1 | 1 | 1 | 1 | 1 |
| LCPAC202 | Pitho-like |  | 26 | 5141 | 72684 | 443964 | 354 | 1 | 2 | 0 | 1 | 2 | 1 | 1 | 1 | 1 | 1 | 1 | 1 | 1 | 0 | 1 |
| LCPAC302 | Pitho-like |  | 30 | 5274 | 20428 | 290561 | 294 | 0 | 1 | 0 | 1 | 1 | 0 | 0 | 1 | 1 | 1 | 1 | 0 | 1 | 0 | 0 |
| LCPAC304 | Pitho-like |  | 12 | 11737 | 173767 | 638759 | 688 | 1 | 1 | 0 | 1 | 1 | 1 | 1 | 1 | 1 | 1 | 0 | 1 | 1 | 1 | 1 |
| LCPAC401 | Pitho-like |  | 11 | 7155 | 114453 | 484752 | 504 | 1 | 1 | 0 | 1 | 1 | 1 | 1 | 1 | 1 | 2 | 1 | 0 | 1 | 1 | 1 |
| LCPAC403 | Pitho-like |  | 6 | 24087 | 117884 | 420388 | 430 | 1 | 1 | 0 | 1 | 1 | 1 | 1 | 1 | 1 | 2 | 0 | 1 | 1 | 1 | 1 |
| LCPAC404 | Pitho-like |  | 10 | 11211 | 84762 | 436585 | 390 | 1 | 1 | 0 | 1 | 1 | 1 | 1 | 1 | 1 | 2 | 0 | 1 | 1 | 1 | 1 |
| LCPAC406 | Pitho-like |  | 10 | 11113 | 75955 | 384297 | 401 | 1 | 1 | 0 | 1 | 1 | 1 | 1 | 1 | 1 | 2 | 1 | 0 | 1 | 1 | 1 |
| LCDPAC01 | Pitho-like |  | 21 | 5383 | 31931 | 282320 | 282 | 1 | 1 | 0 | 1 | 1 | 1 | 1 | 1 | 1 | 0 | 1 | 0 | 1 | 1 | 0 |
| LCDPAC02 | Pitho-like |  | 9 | 6786 | 90916 | 367310 | 390 | 1 | 1 | 0 | 1 | 1 | 1 | 1 | 0 | 0 | 1 | 1 | 0 | 1 | 0 | 0 |
| LCMAC101 | Marseille-like |  | 7 | 15190 | 393561 | 763048 | 793 | 1 | 1 | 1 | 1 | 1 | 1 | 1 | 1 | 1 | 1 | 1 | 1 | 1 | 1 | 1 |
| LCMAC102 | Marseille-like |  | 1 | 395459 | 395459 | 395459 | 465 | 1 | 1 | 1 | 1 | 1 | 1 | 1 | 1 | 1 | 1 | 1 | 1 | 1 | 1 | 1 |
| LCMAC103 | Marseille-like |  | 9 | 14346 | 69824 | 389984 | 427 | 1 | 1 | 1 | 1 | 1 | 1 | 1 | 1 | 0 | 1 | 0 | 1 | 1 | 1 | 0 |
| LCMAC201 | Marseille-like |  | 25 | 6728 | 57873 | 565697 | 566 | 1 | 1 | 1 | 1 | 1 | 0 | 1 | 1 | 0 | 1 | 2 | 1 | 1 | 1 | 1 |
| LCMAC202 | Marseille-like |  | 19 | 6906 | 153726 | 705352 | 672 | 1 | 1 | 2 | 1 | 1 | 1 | 1 | 1 | 0 | 1 | 2 | 1 | 1 | 1 | 1 |
| LCIVAC01 | Iridovirus-like |  | 19 | 5375 | 17223 | 198495 | 222 | 0 | 1 | 0 | 1 | 1 | 1 | 1 | 0 | 1 | 0 | 1 | 1 | 1 | 0 | 1 |
| LCMiAC01 | Mimivirus-like |  | 18 | 8458 | 85120 | 672112 | 571 | 6 | 1 | 1 | 1 | 1 | 1 | 1 | 1 | 1 | 1 | 1 | 1 | 1 | 1 | 0 |
| LCMiAC02 | Mimivirus-like |  | 21 | 8237 | 131456 | 642939 | 583 | 6 | 1 | 2 | 1 | 1 | 2 | 1 | 2 | 1 | 1 | 1 | 1 | 1 | 1 | 1 |
| Cedratvirus A11 |  |  |  |  |  | 589068 | 574 | 1 | 1 | 0 | 1 | 1 | 1 | 1 | 1 | 1 | 1 | 1 | 1 | 1 | 1 | 1 |
| Orpheovirus IHUMI LCC2 |  |  |  |  |  | 1473573 | 1199 | 1 | 1 | 0 | 1 | 1 | 1 | 1 | 1 | 1 | 1 | 1 | 1 | 1 | 1 | 1 |
| Pithovirus sibericum |  |  |  |  |  | 610033 | 425 | 1 | 1 | 0 | 1 | 1 | 1 | 1 | 1 | 1 | 1 | 1 | 1 | 1 | 1 | 1 |
| Marseillevirus |  |  |  |  |  | 369360 | 403 | 1 | 1 | 1 | 1 | 1 | 1 | 1 | 1 | 1 | 1 | 1 | 1 | 1 | 1 | 1 |
| Diadromus pulchellus ascovirus 4a |  |  |  |  |  | 119343 | 119 | 1 | 1 | 1 | 1 | 1 | 1 | 0 | 1 | 1 | 1 | 1 | 1 | 0 | 0 | 1 |
| Heliothis virescens ascovirus 3e |  |  |  |  |  | 186262 | 180 | 1 | 1 | 1 | 1 | 1 | 1 | 0 | 1 | 1 | 1 | 1 | 1 | 0 | 1 | 1 |
| Lymphocystis disease virus |  |  |  |  |  | 186250 | 239 | 1 | 1 | 1 | 1 | 1 | 1 | 1 | 1 | 1 | 1 | 1 | 1 | 0 | 1 | 0 |
| Frog virus 3 |  |  |  |  |  | 105903 | 99 | 1 | 1 | 1 | 1 | 1 | 1 | 0 | 1 | 1 | 1 | 1 | 1 | 0 | 1 | 0 |
| Wiseana iridescent virus |  |  |  |  |  | 205791 | 193 | 1 | 1 | 1 | 1 | 1 | 1 | 1 | 1 | 1 | 1 | 1 | 1 | 1 | 1 | 1 |
| Cafeteria roenbergensis virus BV PW1 |  |  |  |  |  | 617453 | 544 | 3 | 1 | 1 | 1 | 1 | 1 | 1 | 1 | 0 | 1 | 1 | 0 | 1 | 1 | 1 |
| Acanthamoeba polyphaga mimivirus |  |  |  |  |  | 1181549 | 979 | 4 | 1 | 2 | 1 | 1 | 1 | 1 | 1 | 1 | 1 | 1 | 0 | 1 | 1 | 1 |
| Klosneuvirus KNV1 |  |  |  |  |  | 1573084 | 1545 | 7 | 1 | 3 | 1 | 1 | 2 | 1 | 1 | 1 | 1 | 3 | 2 | 1 | 2 | 1 |

Data on representative complete NCLDV genomes from the relevant families are included for comparison.
<sup>a)</sup> MCP, NCLDV major capsid protein (NCVOG0022); DNAp, DNA polymerase family B, elongation subunit (NCVOG0038); ATP, A32-like packaging ATPase (NCVOG0249); RNApA, DNA-directed RNA polymerase subunit alpha (NCVOG0274); RNApB, DNA-directed RNA polymerase subunit beta (NCVOG0271); D5hel, D5-like helicase-primase (NCVOG0023); A18hel, A18-like helicase (NCVOG0076); VLTF3, Poxvirus Late Transcription Factor VLTF3 (NCVOG0262); VLTF2, A1L transcription factor/late TF VLTF-2 (NCVOG1164); RNAp5, DNA-directed RNA polymerase subunit 5 (NCVOG0273); Erv1, Erv1/Alr family disulfide (thiol) oxidoreductase (NCVOG0052); RNAlig, RNA ligase (NCVOG1088); TopoII, DNA topoisomerase II (NCVOG0037); FLAP, Flap endonuclease (NCVOG1060); TFIIIB, transcription initiation factor IIB (NCVOG1127).

Differential coverage binning was performed first, resulting in 29 genomic bins. Initial quality assessment showed that most of the bins were inflated and fragmented, containing many short contigs (<5kb), which were difficult to classify as contamination or *bona fide* NCLDV sequences, and some bins were likely to contain sequences from more than one viral genome, judged by the presence of marker genes belonging to different families of the NCLDV (Additional file 1, Figures S19-S20). The more contigs a bin contains, the higher the risk is that some of these could be contaminants that bin together because of similar nucleotide composition and read coverage. Therefore, sequence read profiling followed by co-assembly binning was performed in an attempt to increase the size of the contigs and thus obtain additional information for binning and bin refinement. For most of the bins, the co-assembly led to a decrease in the number of contigs, without losing completeness or even improving it (Additional file 1, Table S6).

A key issue with metagenomic binning is whether contigs are binned together because they belong to the same genome, or rather because they simply display a similar nucleotide composition and read coverage. In general, contigs were retained if they contained at least one gene with BLASTP top hits to NCLDV proteins. Some contigs encoded proteins with only bacterial, archaeal, and/or eukaryotic BLASTP top hits, and because the larger NCLDV genomes contain islands enriched in genes of bacterial origin (43, 49), it was unclear which sequences could potentially be contaminants. A combination of gene content, coverage and composition information was used to identify potential contaminating sequences. Contigs shorter than 5 kb were also discarded because they generally do not contain enough information to reliably establish their origin, but this strict filtering also means that the size of the genomes could be underestimated and some genomic information lost. Reassuringly, no traces of ribosomal RNA or ribosomal protein genes were identified in any of the NCLDV genome bins, which would have been a clear case of contaminating cellular sequences. Altogether, of the 336 contigs in the 23 final genome bins, 243 (72%) could be confidently assigned to NCLDV on the basis of the presence of at least one NCLDV-specific gene.

The content of the 23 NCLDV-related bins was analyzed in more depth (Table 1). The bins included from 1 to 30 contigs, with the total length of non-overlapping sequences varying from about 200 to more than 750 kilobases (kb), suggesting that some might contain (nearly) complete NCLDV genomes although it is difficult to make any definitive conclusions on completeness from length alone because the genome size of even closely related NCLDV can vary substantially. A much more reliable approach is to assess the representation of core genes that are expected to be conserved in (nearly) all NCLDV. The translated protein sequences from the 23 bins were searched for homologs of conserved NCLDV genes using PSI-BLAST, with profiles of the NCVOGs employed as queries ((28); see Additional File 2 for protein annotation). Of the 23 bins, in 14 (nearly) complete sets of the core NCLDV genes were identified (Table 1) suggesting that these bins contained (nearly) complete genomes of putative new viruses (hereafter, LCV, Loki’s Castle Viruses). Notably, the Pithovirus-like LCV lack the packaging ATPase of the FtsK family that is encoded in all other NCLDV genomes but not in the available Pithovirus genomes. Several bins contained more than one copy of certain conserved genes. Some of these could represent actual paralogs but, given that duplication of most of these conserved genes (e.g. DNA Polymerase in Bin LCPAC202 or RNA polymerase B subunit in Bins LCPAC201 and LCPAC202) is unprecedented among NCLDV, it appears likely that several bins are heterogeneous, each containing sequences from two closely related virus genomes.

With all the caution due because of the lack of fully assembled virus genomes, the range of the apparent genomes sizes of the Pitho-like and Marseille-like LCV is notable (Table 1). The characteristic size of the genomes in the family ”Pithoviridae” is about 600 kb (17-19) but, among the Pitho-like LCV, only one, LCPAC304, reached and even exceeded that size. The rest of the LCV genomes are substantially smaller, and although some are likely to be incomplete, given that certain core genes are missing, others, such as LCPAC104, with the total length of contigs at only 218 kb, encompass all the core genes (Table 1).

The typical genome size in the family *Marseilleviridae* is between 350 and 400 kb (22) but among the LCV, genomes of two putative Marseille-like viruses, LCMAC101 and LCMAC202, appear to exceed 700 kb, well into the giant virus range. Although LCMAC202 contains two uncharacteristic duplications of core genes, raising the possibility of heterogeneity, LCMAC101 contains all core genes in a single copy, and thus, appears to be an actual giant virus. Thus, the family *Marseilleviridae* seems to be joining the NCLDV families that evolved virus gigantism.

A concatenation of the three most highly conserved proteins, namely, NCLDV major capsid protein (MCP), DNA polymerase (DNAP), and A18-like helicase (A18Hel), was used for phylogenetic analysis (see Methods for details). Among the putative new NCLDV, 15 cluster with Pithoviruses (Figure 3). These new representatives greatly expand the scope of the family ”Pithoviridae”. Indeed, 8 of the 15 form a putative (weakly supported) clade that is the sister group of all currently known ”Pithoviridae” (Pithovirus, Cedratvirus and Orpheovirus), 5 more comprise a deeper clade, and LCDPAC02 represents the deepest lineage of the Pitho-like viruses (Figure 3). Additionally, 5 of the putative new NCLDV are affiliated with the family *Marseilleviridae*, and similarly to the case of Pitho-like viruses, two of these comprise the deepest branch in the Marseille-like subtree (although the monophyly of this subtree is weakly supported) (Figure 3). Another LCV represents a distinct lineage within the family *Iridoviridae* (Figure 3). The topologies of the phylogenetic trees for individual conserved NLCDV genes were mostly compatible with these affinities of the putative new viruses Additional File 3). Taken together, these findings substantially expand the Pitho-Irido-Marseille (PIM) clade of the NCLDV, and the inclusion of the LCV in the phylogeny confidently reaffirms the previously observed monophyly of this branch (Figure 3). Finally, two LCV belong to the Klosneuvirus branch (putative subfamily “Klosneuvirinae”) within the family *Mimiviridae* (Figure 3, inset).

**Figure 3.**
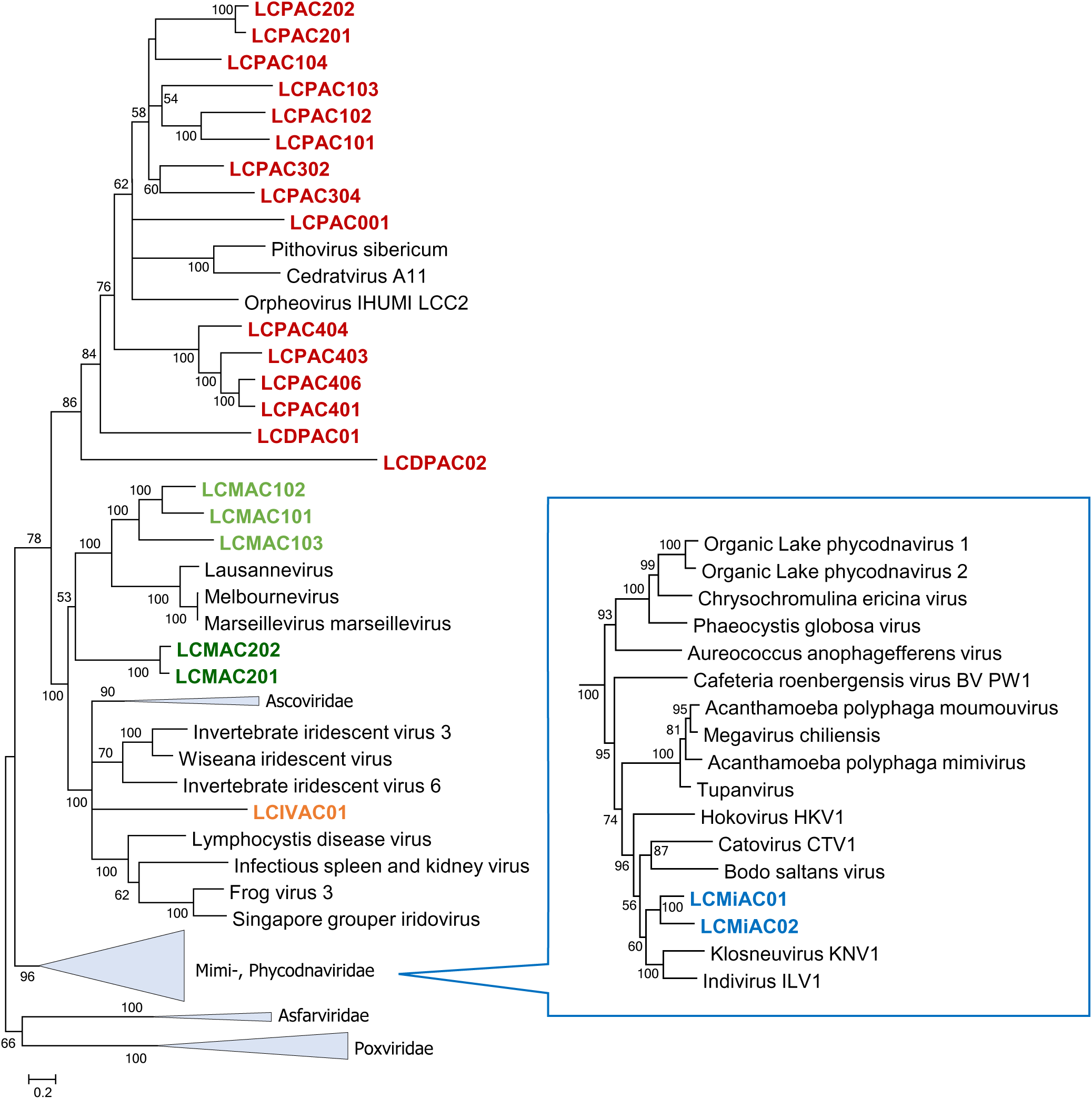
Phylogenetic tree of three concatenated, universally conserved NCLDV proteins: DNA polymerase, major capsid protein, and A18-like helicase. Support values were obtained using 100 bootstrap replications; branches with support less than 50% were collapsed. Scale bars represent the number of amino acid (aa) substitutions per site. The inset shows the *Mimiviridae* branch. Triangles show collapsed branches. The LCV sequences are color-coded as follows: red, Pitho-like; green, Marseille-like (a deep branch shown in dark green); orange, Irido-like; blue, Mimi (Klosneu)-like.

### Translation system components encoded by Loki’s Castle viruses

Similar to other NCLDV with giant and large genomes, the LCV show a patchy distribution of genes coding for translation system components. Such genes were identified in 11 of the 23 bins (Table 2; Additional File 2). None of the putative new viruses has a (near) complete set of translation-related genes (minus the ribosome) as observed in Klosneuviruses (46) or Tupanviruses (98). Nevertheless, several of the putative Pitho-like viruses encode multiple translation-related proteins, e.g. Bin LCMAC202 that encompasses 6 aminoacyl-tRNA synthetases (aaRS) and 6 translation factors or Bin LCMAC201, with 4 aaRS and 5 translation factors (Table 2). Additionally, 12 of the 23 bins encode predicted tRNAs, up to 22 in Bin LCMAC202 (Table 2).

**Table 2.** Translation-related proteins and tRNAs in Loki’s Castle NCLDV

| bin name | AlaS<sup>a</sup> | AsnS | GRS1 | GlnS | HisS | IleS | MetS | ProS | Pth2 | RLI1 | ThrS | TrpS | TyrS | eIF1 | eIF1a | eIF2a | eIF2b | eIF2g | eIF4e | eIF5b | eRF1 | tRNA |
| --- | --- | --- | --- | --- | --- | --- | --- | --- | --- | --- | --- | --- | --- | --- | --- | --- | --- | --- | --- | --- | --- | --- |
| LCPAC001 |  | 1 |  |  |  |  | 1 |  |  |  |  |  |  |  |  |  |  |  |  |  |  | 5 |
| LCPAC101 |  |  | 1 |  |  |  |  |  |  |  |  |  |  |  |  |  |  |  |  |  |  | 2 |
| LCPAC102 |  |  |  |  |  |  |  |  |  |  |  |  |  |  |  |  |  |  |  |  |  | 3 |
| LCPAC103 |  |  |  |  |  |  |  |  |  |  |  |  |  |  |  |  |  |  |  |  |  |  |
| LCPAC104 |  |  |  |  |  |  |  |  |  |  |  |  |  |  |  |  |  |  |  |  |  | 4 |
| LCPAC201 |  |  |  |  |  |  |  |  |  |  |  |  |  |  |  |  |  |  |  |  |  |  |
| LCPAC202 |  |  |  |  |  |  |  |  |  |  |  |  |  |  |  |  |  |  |  |  |  |  |
| LCPAC302 |  |  |  |  |  |  |  |  |  | 1 |  |  |  |  |  |  |  |  |  |  |  |  |
| LCPAC304 |  | 1 |  |  |  |  |  |  | 1 | 1 |  | 1 |  |  |  |  | 1 |  |  |  |  | 21 |
| LCPAC401 |  |  |  |  |  |  |  |  |  |  |  |  |  |  |  |  |  |  |  |  |  |  |
| LCPAC403 |  |  |  |  |  |  |  |  |  |  |  |  |  |  |  |  |  |  |  |  |  |  |
| LCPAC404 |  |  |  |  |  |  |  |  |  |  |  |  |  |  |  |  | 1 |  |  |  |  |  |
| LCPAC406 |  |  |  |  |  |  |  |  |  |  |  |  |  |  |  |  |  |  |  |  |  |  |
| LCDPAC01 |  |  |  |  |  |  |  |  |  |  |  |  |  |  |  |  |  |  |  |  |  |  |
| LCDPAC02 |  |  |  |  |  |  |  |  |  |  |  |  |  |  |  |  |  |  |  |  |  |  |
| LCMAC101 |  | 3 |  |  |  |  |  |  | 1 |  |  |  |  |  |  |  |  |  |  |  |  | 8 |
| LCMAC102 |  |  |  |  |  |  |  |  |  |  |  |  |  |  |  |  |  |  |  |  |  | 3 |
| LCMAC103 | 1 |  |  |  |  |  |  |  |  |  |  |  |  |  |  | 1 | 1 | 1 | 1 | 1 | 1 | 17 |
| LCMAC201 |  | 1 |  | 1 |  |  | 1 |  |  |  |  |  | 1 |  |  | 1 | 2 | 1 |  |  |  | 11 |
| LCMAC202 | 1 | 2 |  |  |  |  |  | 1 |  |  | 1 |  | 1 | 1 |  | 1 | 2 | 1 |  |  | 1 | 26 |
| LCIVAC01 |  |  |  |  |  |  |  |  |  |  |  |  |  |  |  |  |  |  |  |  |  |  |
| LCMiAC01 |  |  |  |  | 1 | 1 |  |  |  |  | 1 |  | 1 |  | 1 |  |  |  | 1 |  |  | 18 |
| LCMiAC02 |  |  |  |  |  |  |  |  |  |  |  |  |  |  |  |  |  |  | 2 |  | 1 | 2 |
| Pithovirus sibericum |  |  |  |  |  |  |  |  |  |  |  |  |  |  |  |  |  |  |  |  |  |  |
| Cedratvirus_A11 |  |  |  |  |  |  |  |  |  |  |  |  |  |  |  |  |  |  |  |  |  |  |
| Orpheovirus |  | 1 | 1 |  | 1 | 1 |  |  |  |  |  |  | 1 | 1 |  |  |  |  |  |  | 1 |  |
| Marseillevirus |  |  |  |  |  |  |  |  |  |  |  |  |  | 1 |  |  | 1 |  |  |  | 1 |  |
| Klosneuvirus_KNV1 | 1 | 1 | 1 | 2 | 1 | 1 | 1 | 1 | 3 | 1 | 1 | 1 | 1 | 1 | 1 | 1 | 2 | 1 | 1 |  | 1 | 25 |
| mimivirus |  | 1 |  |  |  | 1 | 1 |  |  |  |  |  | 1 | 1 |  |  |  |  | 1 |  | 2 | 6 |
| Tupanvirus | 1 | 1 | 1 | 2 | 1 | 1 | 1 | 1 |  |  | 1 | 1 | 1 | 1 | 1 | 1 | 2 | 1 | 2 |  | 1 |  |
| C. roenbergensis virus |  |  |  |  |  | 1 |  |  |  |  |  |  |  | 1 | 1 | 1 | 3 | 1 | 1 | 1 |  | 16 |

<sup>a</sup> translation-related proteins are abbreviated as follows: AlaS, Alanyl-tRNA synthetase; AsnS, Aspartyl/asparaginyl-tRNA synthetase; GlnS, Glutamyl- or glutaminyln-tRNA synthetase; GRS1, Glycyl-tRNA synthetase (class II) ; HisS, Histidyl-tRNA synthetase; IleS, Isoleucyl-tRNA synthetase; MetS, Methionyl-tRNA synthetase; ProS, Prolyl-tRNA synthetase; ThrS, Threonyl-tRNA synthetase; TrpS, Tryptophanyl-tRNA synthetase; TyrS, Tyrosyl-tRNA synthetase; Pth2, Peptidyl-tRNA hydrolase ; eIF1, Translation initiation factor 1 (eIF-1/SUI1); eIF1a, Translation initiation factor 1A/IF-1; eIF2a, Translation initiation factor 2, alpha subunit (eIF-2alpha) ; eIF2b, Translation initiation factor 2, beta subunit (eIF-2beta)/eIF-5 N-terminal domain ; eIF2g, Translation initiation factor 2, gamma subunit (eIF-2gamma; GTPase) ; eIF4e, Translation initiation factor 4E (eIF-4E); eIF5b, Translation initiation factor IF-2 (Initiation Factor 2 (IF2)/ eukaryotic Initiation Factor 5B (eIF5B) family; IF2/eIF5B); eRF1, Peptide chain release factor 1 (eRF1) ; RLI1, Translation initiation factor RLI1
Data for completely sequenced representatives of the relevant NCLDV families are included for comparison.

Given the special status of the translation system components in the discussions of the NCLDV evolution, we constructed phylogenies for all these genes including the LCV and all other NCLDV. The results of this phylogenetic analysis (Figure 4 and Additional File 3) reveal complex evolutionary trends some of which that have not been apparent in previous analyses of the NCLDV evolution. First, in most cases when multiple LCV encompass genes for homologous translation system components, phylogenetic analysis demonstrates polyphyly of these genes. Notable examples include translation initiation factor eIF2b, aspartyl/asparaginyl-tRNA synthetase (AsnS), tyrosyl-tRNA synthetase (TyrS) and methionyl-tRNA synthetase (MetS; Figure 4). Thus, the eIF2b tree includes 3 unrelated LCV branches one of which, not unexpectedly, clusters with homologs from Marseilleviruses and Mimiviruses, another one is affiliated with two Klosneuviruses, and the third one appears to have an independent eukaryotic origin (Figure 4a). The AsnS tree includes a group of LCV that clusters within a mixed bacterial and archaeal branch that also includes two other NCLDV, namely, Hokovirus of the Klosneuvirus group and a phycodnavirus. Another LCV AsnS belongs to a group of apparent eukaryotic origin and one, finally, belongs to a primarily archaeal clade (Figure 4b and Additional File 3). Of the 3 TyrS found in LCV, two cluster with the homologs from Klosneuviruses within a branch of apparent eukaryotic origin, and the third one in another part of the same branch where it groups with the Orpheovirus TyrS; notably, the same branch includes homologs from pandoraviruses (Figure 4c). Of the two MetS, one groups with homologs from Klosneuviruses whereas the other one appears to be of an independent eukaryotic origin (Figure 4d). These observations are compatible with the previous conclusions on multiple, parallel acquisitions of genes for translation system components by different groups of NCLDV (primarily, giant viruses but, to a lesser extent, also those with smaller genomes), apparently, under evolutionary pressure for modulation of host translation that remains to be studied experimentally.

**Figure 4.**
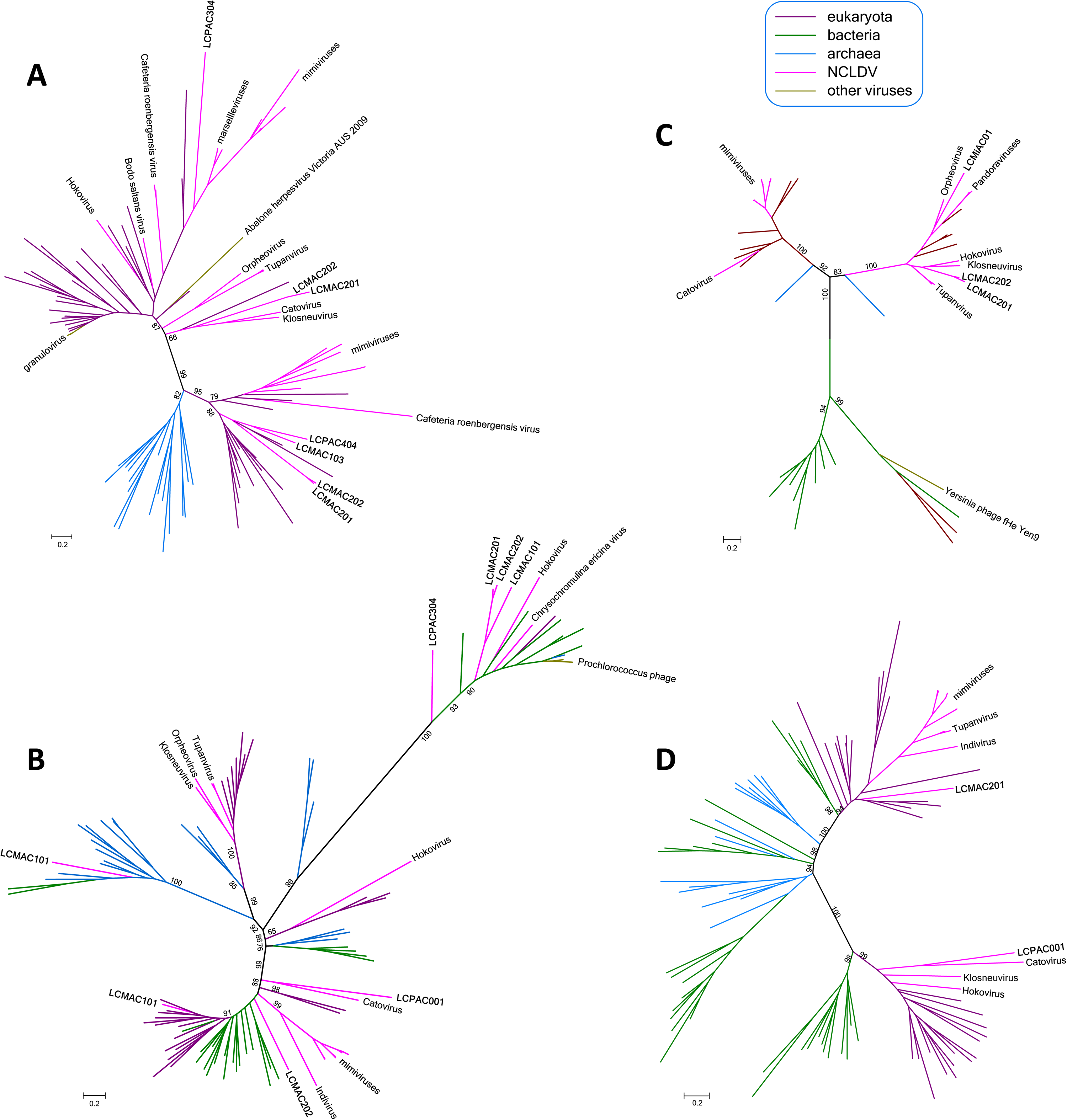
Phylogenies of selected translation system components encoded by Loki’s Castle viruses. A, translation initiation factor eIF2b B, aspartyl/asparaginyl-tRNA synthetase, AsnS C, tyrosyl-tRNA synthetase, TyrS, D, methionyl-tRNA synthetase, MetS. All branches are color-coded according to taxonomic affinity (see Additional File 3 for the full trees). The numbers at the internal branches indicate local likelihood-based support (percentage points).

Another clear trend among the translation-related genes of the Pitho-like LCV is the affinity of several of them with homologs from Klosneuviruses and, in some cases, Mimiviruses. All 4 examples mentioned about include genes of this provenance, and additional cases are GlyS, IleS, ProS, peptidyl-tRNA hydrolase, translation factors eIF1a and eIF2a, and peptide chain release factor eRF1 (Additional File 3). Given that the LCV set includes two Klosneuvirus-like bins, in addition to the Pitho-like ones, these observations imply extensive gene exchange between distinct NCLDV in the habitats from which these viruses originate. Klosneuviruses that are conspicuously rich in translation-related genes might serve as the main donors.

### Gene content analysis of the Loki’s Castle viruses

Given that the addition of the LCV has greatly expanded the family *Marseilleviridae* and the Pithovirus group, and reaffirmed the monophyly of the PIM branch of NCLDV, we constructed, analyzed and annotated clusters of putative orthologous genes for this group of viruses as well as an automatically generated version of clusters of homologous genes for all NCLDV (ftp://ftp.ncbi.nih.gov/pub/yutinn/Loki_Castle_NCLDV_2018/NCLDV_clusters/). Altogether, 8066 NCLDV gene clusters were identified of which a substantial majority were family-specific. Nevertheless, almost 200 clusters were found to be shared between Pithoviridae and *Marseilleviridae* families (Figure 5). The numbers of genes shared by each of these families with *Iridoviridae* were much smaller, conceivably, because of the small genome size of iridoviruses that could have undergone reductive evolution (Figure 5). Conversely, there was considerable overlap between the PIM group gene clusters and those of mimiviruses, presumably, due to the large genome sizes of the mimiviruses, but potentially reflecting also substantial horizontal gene flow between mimiviruses and pitho- and marseilleviruses (Figure 5). Only 13 genes comprised a genomic signature of the PIM group, that is, genes that were shared by its three constituent families, to the exclusion of the rest of the NCLDV.

**Figure 5.**
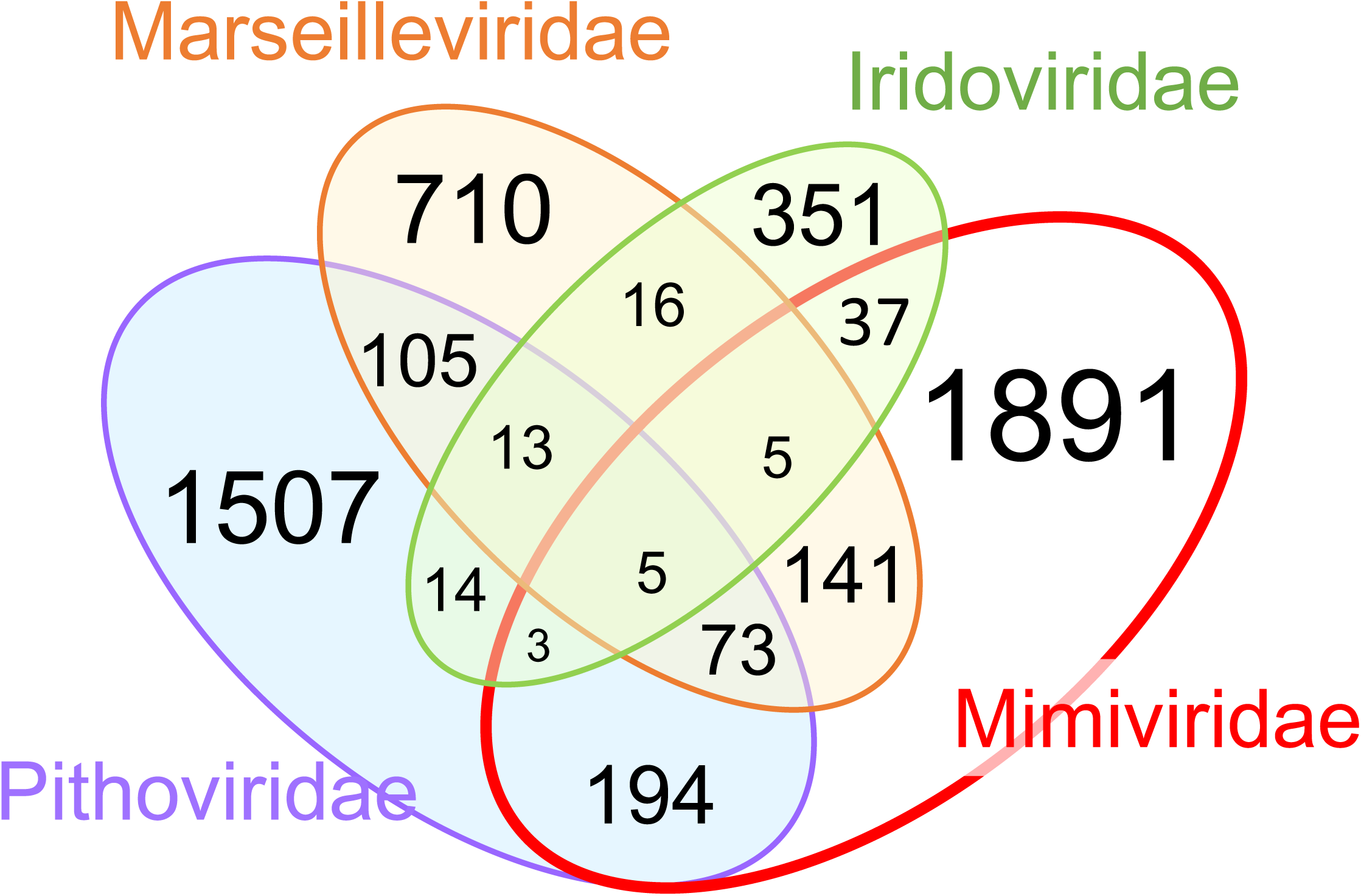
Shared and unique genes in four NCLDV families that include Loki’s Castle viruses. The numbers correspond to NCLDV clusters that contain at least one protein from Mimi-, Marseille-, Pitho, and -Iridoviridae, but are absent from other NCLDV families.

To further explore the relationships between the gene repertoires of the PIM group and other NCLDV, we constructed a neighbor-joining tree from the data on gene presence-absence (ftp://ftp.ncbi.nih.gov/pub/yutinn/Loki_Castle_NCLDV_2018/NCLDV_clusters/). Notwithstanding the limited gene sharing, the topology of the resulting tree (Figure 6) closely recapitulated the phylogenetic tree of the conserved core genes (Figure 3). In particular, the PIM group appears as a clade in the gene presence-absence tree albeit with a comparatively low support (Figure 6). Thus, despite the paucity of PIM-specific genes and the substantial differences in the genome sizes between the three virus families, gene gain and loss processes within the viral genetic core appear to track the evolution of the universally conserved genes.

**Figure 6.**
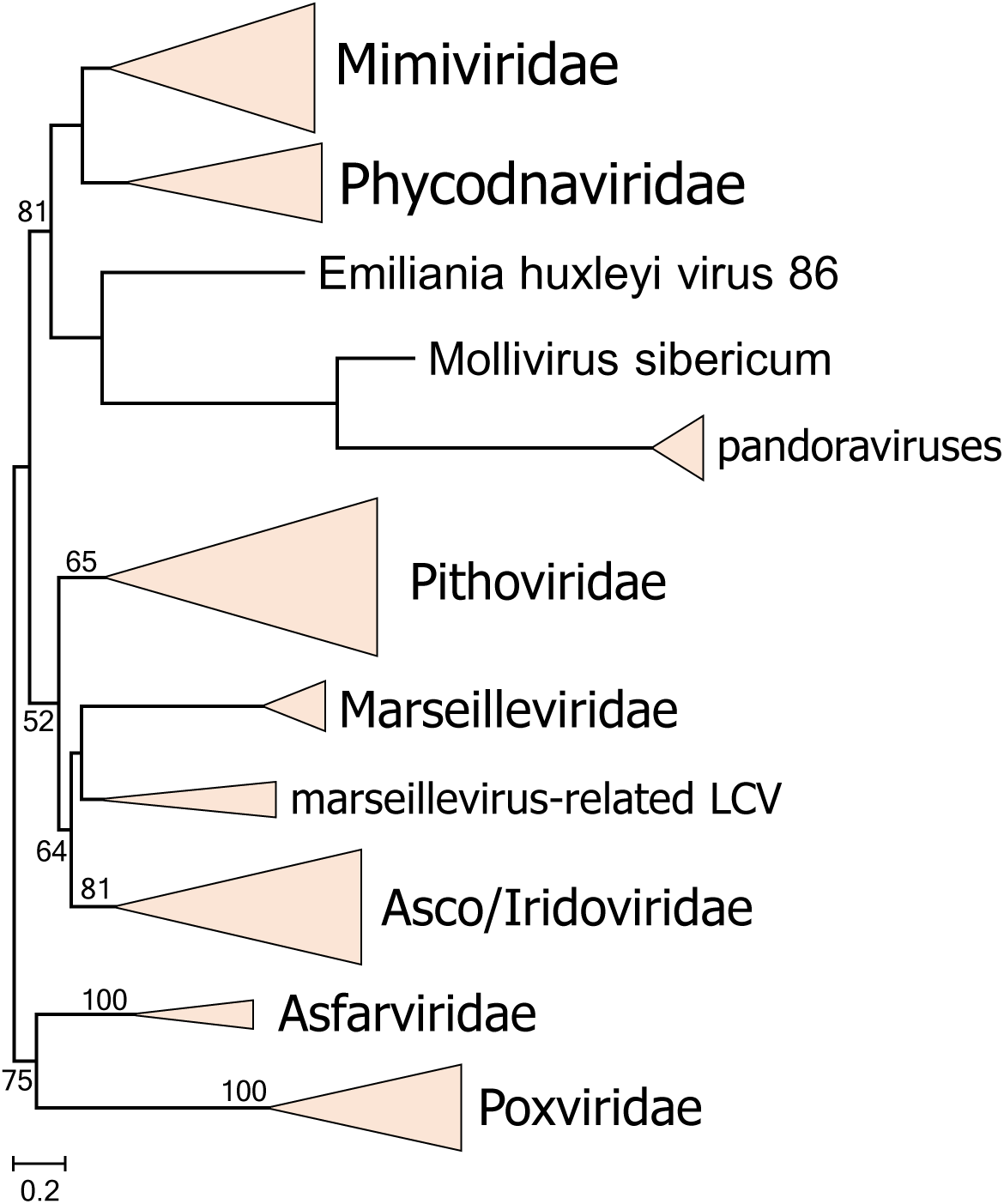
Gene presence-absence tree of the NCLDV including the Loki’s Castle viruses. The Neighbor-Joining dendrogram was reconstructed from the matrix of pairwise distances calculated from binary phyletic patterns of the NCLDV clusters. The numbers at internal branches indicate bootstrap support (percentage points); numbers below 50% are not shown.

The genomes of microbes and large viruses encompass many lineage-specific genes (often denoted ORFans) that, in the course of evolution, are lost and gained by horizontal gene transfer at extremely high rates (99). Therefore, the gene repertoire of a microbial or viral species (notwithstanding the well-known difficulties with the species definition) or group is best characterized by the pangenome, i.e. the entirety of genes represented in all isolates in the group (100-102). Most microbes have “open” pangenomes such that every sequenced genome adds new genes to the pangenome (102, 103). The NCLDV pangenomes could be even wider open, judging from the high percentage of ORFans, especially, in giant viruses (104). Examination of the PIM genes clusters shows that 757 of the 1572 clusters (48%) were unique to the LCV, that is, had no detectable homologs in other members of the group. Taking into account also the 4147 ORFans, the LCV represent the bulk of the PIM group pangenome. Among the NCLDV clusters, 1100 of the 8066 (14%) are LCV-specific. Thus, notwithstanding the limitations of the automated clustering procedure that could miss some distant similarities between proteins, the discovery of the LCV substantially expands not only the pangenome of the PIM group but also the overall NCLDV pangenome.

Annotation of the genes characteristic of (but not necessarily exclusive to) the PIM group reveals numerous, highly diverse functions of either bacterial or eukaryotic provenance as suggested by the taxonomic affiliations of homologs detected in database searches (Additional file 5). For example, a functional group of interest shared by the three families in the PIM group include genes of apparent bacterial origin involved in various DNA repair processes and nucleotide metabolism. The results of phylogenetic analysis of these genes are generally compatible with bacterial origin although many branches are mixed, including also archaea and/or eukaryotes and indicative of horizontal gene transfer (Figure 7). Notably, these trees illustrate the “hidden complexity” of NCLDV evolution whereby homologous genes are independently captured by different groups of viruses. In the trees for the two subunits of the SbcCD nuclease, the PIM group forms a clade but the homologs in mimiviruses appear to be of distinct origin (Figure 7A,B) whereas in the trees for exonuclease V and dNMP kinase, the PMI group itself splits between 3 branches (Figure 7C,D). The latter two trees also contain branches in which different groups of the NCLDV, in particular, marseilleviruses and mimiviruses, are mixed, apparently reflecting genes exchange between distinct viruses infecting the same host, such as amoeba.

**Figure 7.**
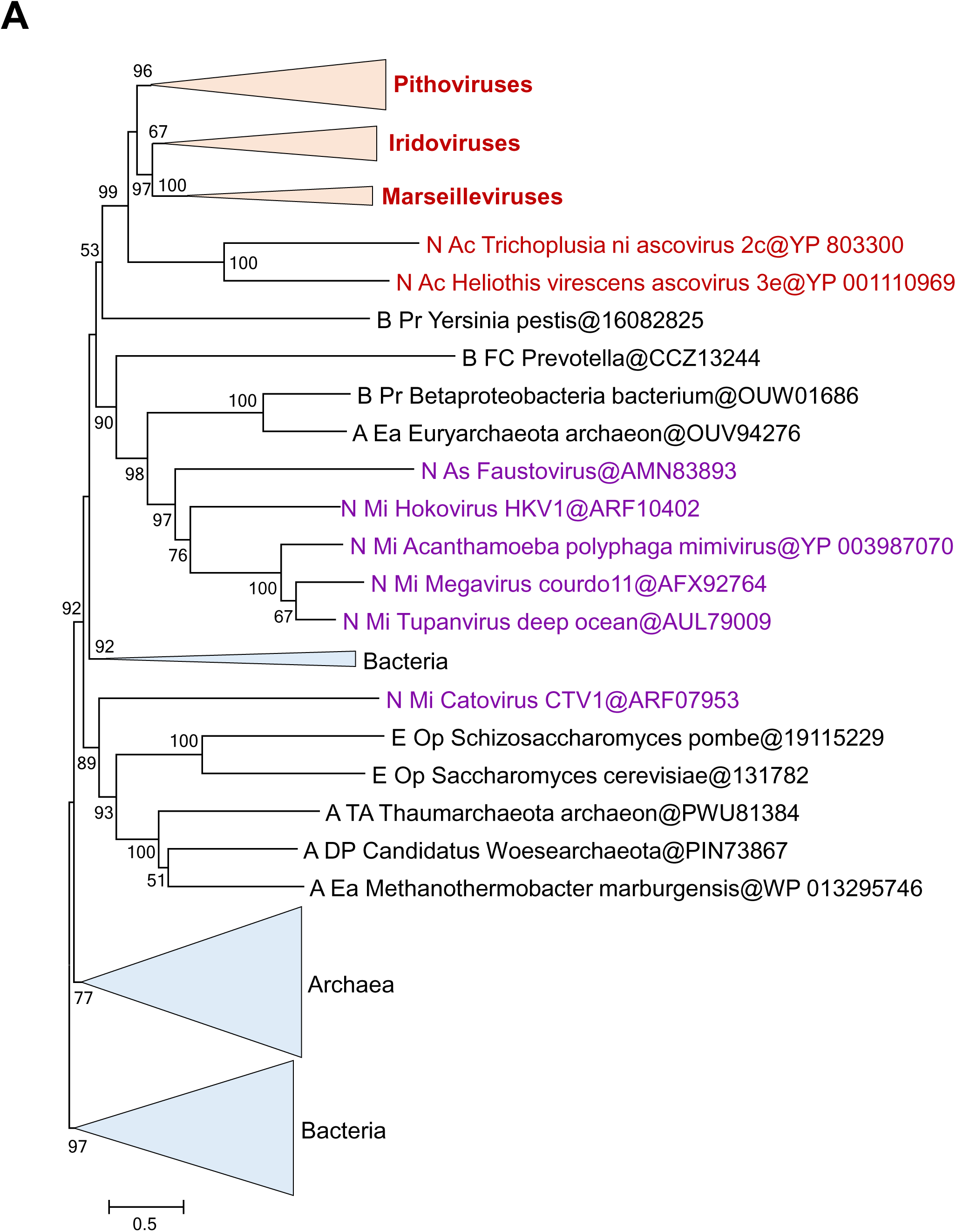
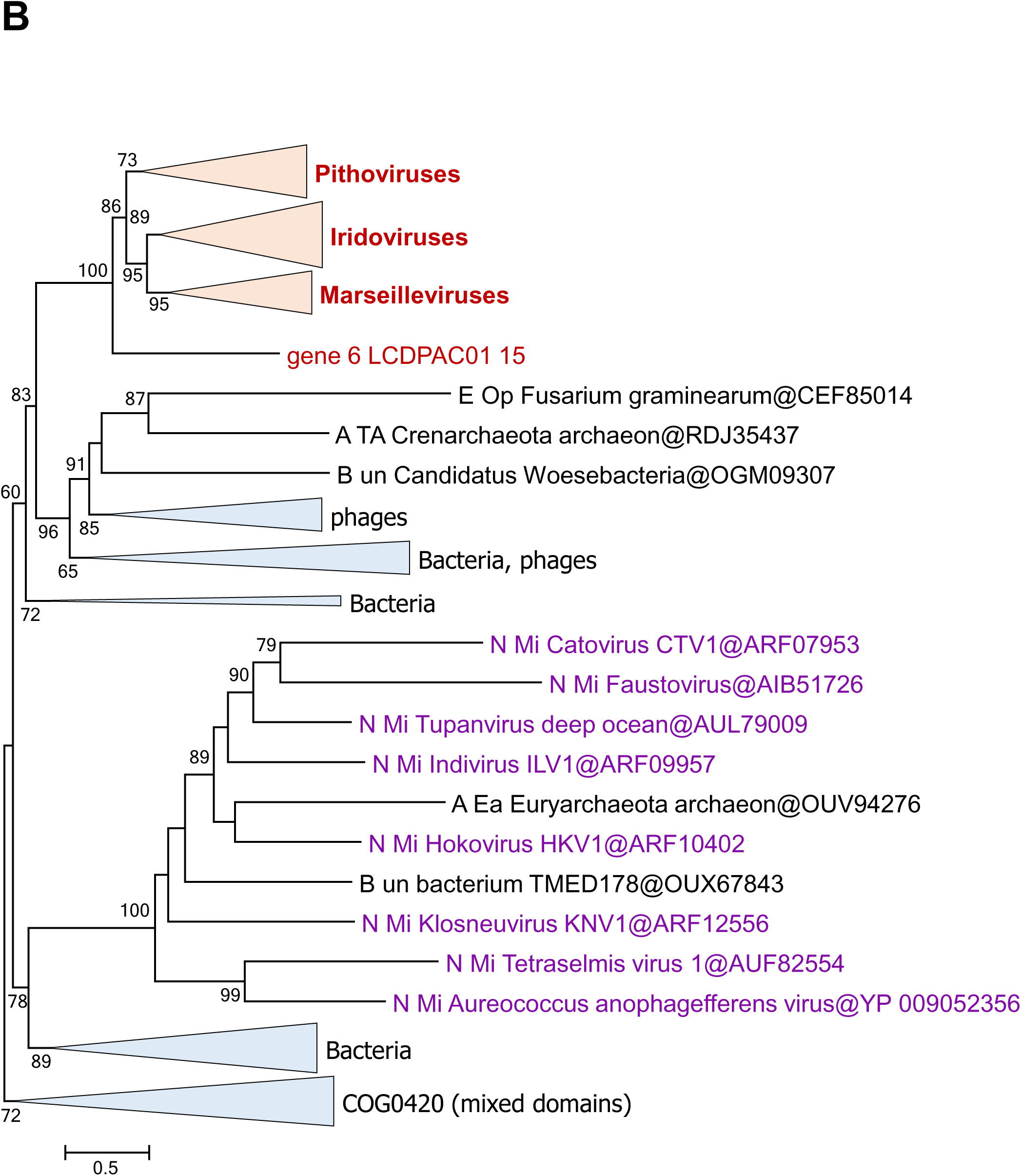
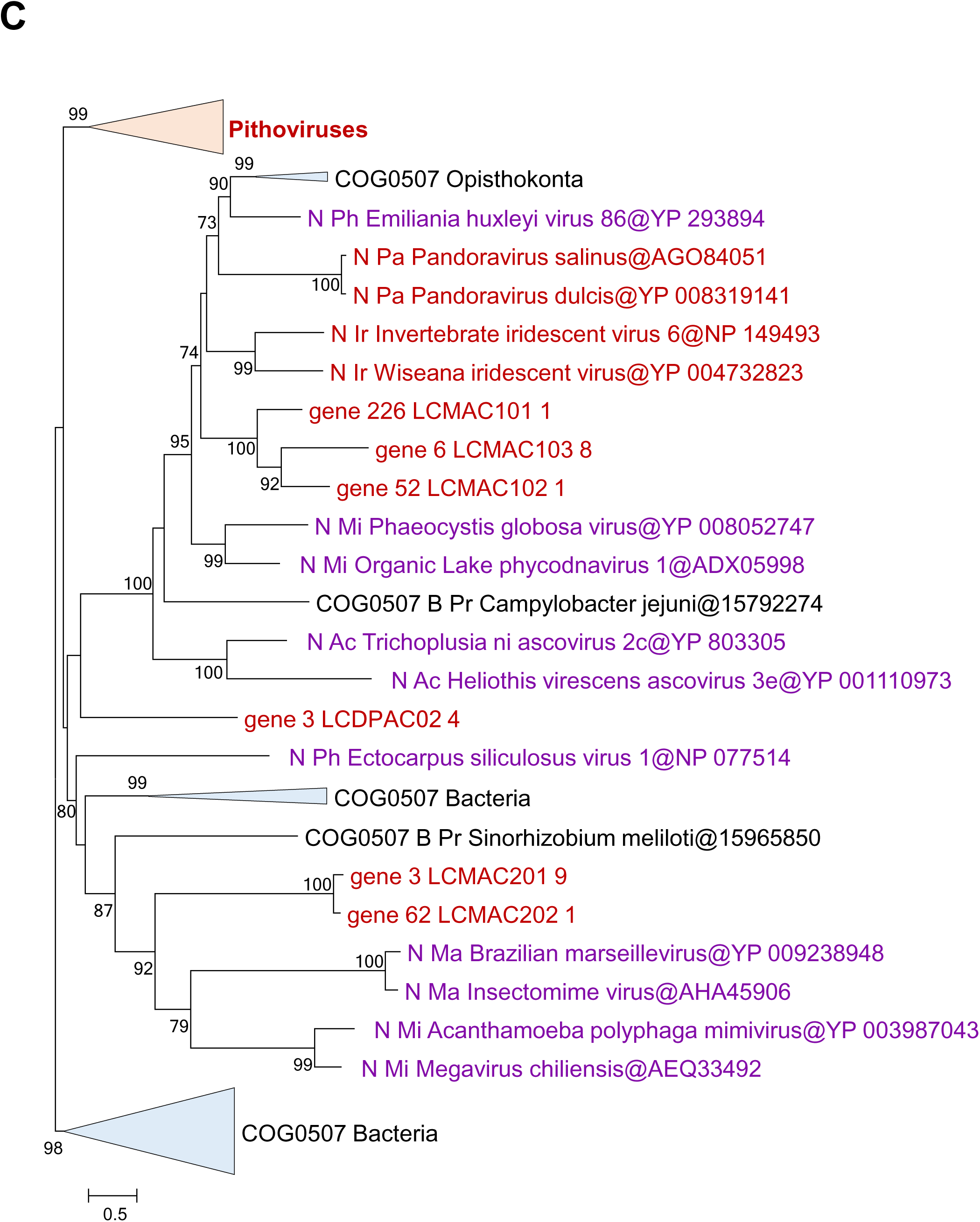
Phylogenies of selected repair and nucleotide metabolism genes of the Pitho-Irido-Marseille virus group including Loki’s Castle viruses. A, SbcCD nuclease, ATPase subunit SbcC B, SbcCD nuclease, nuclease subunit SbcD C, exonuclease V; D, dNMP kinase. The numbers at the internal branches indicate local likelihood-based support (percentage points). Genbank protein IDs, wherever available, are shown after ‘@’. Taxa abbreviations are as follows: A DP, Archaea; DPANN group; A TA, Thaumarchaeota; A Ea, Euryarchaeota; B FC, Bacteroidetes; B Fu, Fusobacteria; B Pr, Proteobacteria; B Te, Firmicutes; B un, unclassified Bacteria; E Op, Opisthokonta; N Pi, ”Pithoviridae”; N Ac, Ascoviridae; N As, Asfarviridae; N Ma, Marseilleviridae; N Mi, Mimiviridae; N Pa, Pandoraviridae; N Ph, Phycodnaviridae; V ds, double-strand DNA viruses.

### Loki’s Castle virophages

Many members of the family *Mimiviridae* are associated with small satellite viruses that became known as virophages (subsequently classified in the family *Lavidaviridae* (105-111). Two virophage-like sequences were retrieved from Loki Castle metagenomes. According to the MCP phylogeny, they form a separate branch within the Sputnik-like group (Figure 8A). This affiliation implies that these virophages are parasites of mimiviruses. Both Loki’s Castle virophages encode the core virophage genes encoding the proteins involved in virion morphogenesis, namely, MCP, minor capsid protein, packaging ATPase, and cysteine protease (Figure 8B and Additional File 2 for protein annotations). Apart from these core genes, however, these virophages differ from Sputnik. In particular, they lack the gene for the primase-helicase fusion protein that is characteristic of Sputnik and its close relatives (112), but each encode a distinct helicase (Figure 8B).

**Figure 8.**
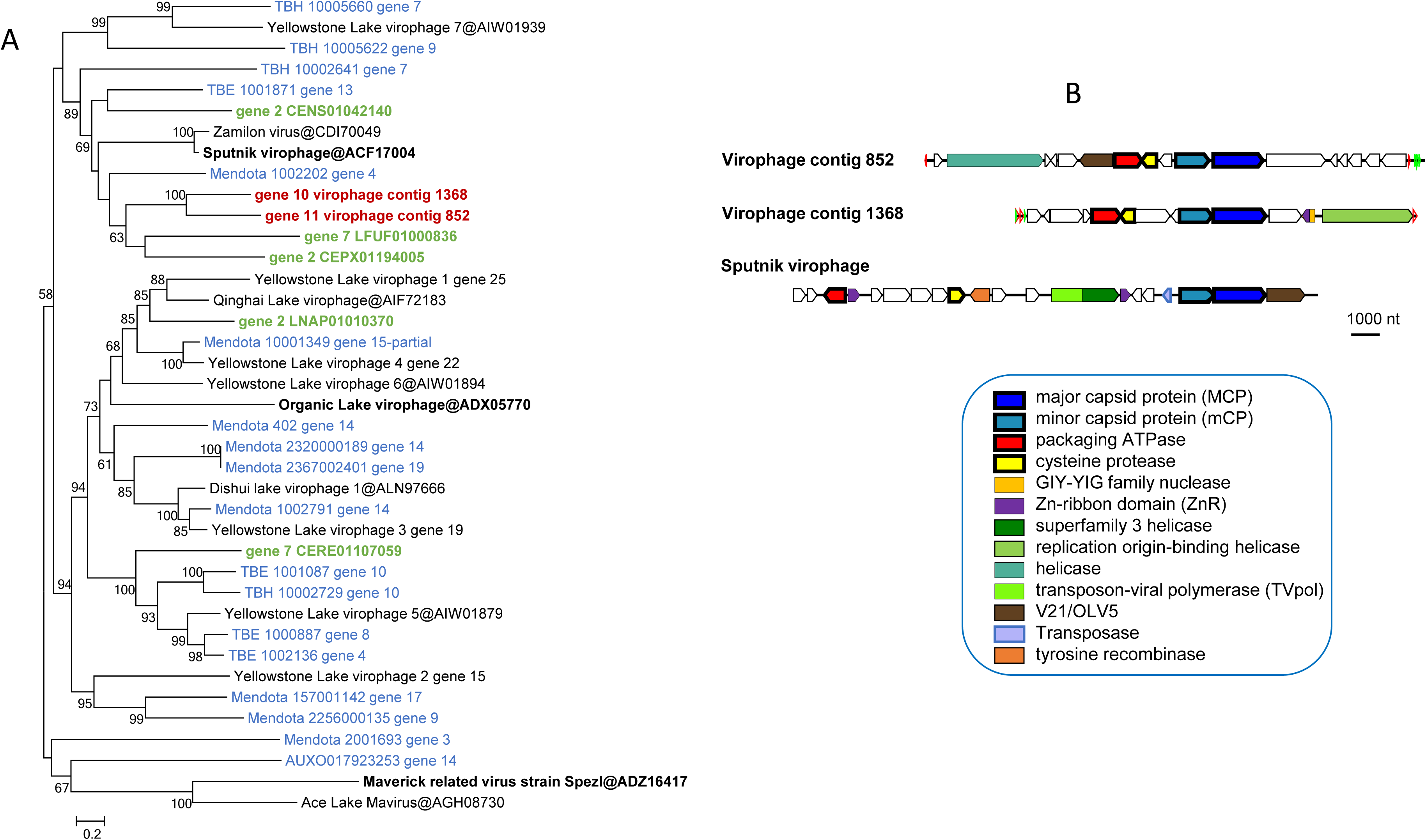
Loki’s Castle virophages. A, Phylogenetic tree of virophage major capsid proteins. Reference virophages from GenBank are marked with black font (the three prototype virophages are shown in bold), environmental virophages shown in blue (128) and green (wgs portion of GenBank). B, Genome maps of Loki’s Castle virophages compared with Sputnik virophage. Green and blue triangles mark direct and inverted repeats. Pentagons with a thick outline represent conserved virophage genes.

### Putative promoter motifs in LCV and Loki’s Castle virophages

To identify possible promoter sequences in the LCV genomes, we searched upstream regions of the predicted LCV genes for recurring motifs using the MEME software (see Methods for details). In most of the bins, we identified a conserved motif similar to the early promoters of poxviruses and mimiviruses (113) (AAAnTGA) that is typically located within 40 to 20 nucleotides upstream of the predicted start codon (for the search results, see: ftp://ftp.ncbi.nih.gov/pub/yutinn/Loki_Castle_NCLDV_2018/meme_motif_search/). To assess possible bin contamination, we calculated frequencies of the conserved motifs per contig, for Marseillevirus-like and Mimivirus-like bins. None of the contigs showed significantly reduced frequency of the conserved motif (Additional file 7), supporting the virus origin of all the contigs.

Notably, the LCV virophage genomes also contain a conserved AT-rich motif upstream of each gene which is likely to correspond to the late promoter of their hosts, similarly to the case of the Sputnik virophage that carries late mimivirus promoters (114). However, the genomes of the two putative Klosneuviruses LCMiAC01 nor LCMiAC02 that are represented among the LCV do not contain obvious counterparts to these predicted virophage promoters (Additional file 8). Therefore, it appears most likely that the hosts of these virophages are mimiviruses that are not represented in the LCV sequence set.

Of further interest is the detection of pronounced promoter-like motifs for pitho-like LCV (Additional file 9) and irido-like LCV (Additional file 10). To our knowledge, no conserved promoter motifs have been so far identified for these groups of viruses.

## Discussion

Metagenomics has become the primary means of new virus discovery (51, 52, 115). Metagenomic sequence analysis has greatly expanded many groups of viruses such that the viruses that have been identified earlier by traditional methods have become isolated branches in the overall evolutionary trees in which most of the diversity comes from metagenomic sequences (116-121). The analysis of the Loki’s Castle metagenome reported here has similarly expanded the Pithovirus branch of the NCLDV, and to a somewhat lesser extent, the Marseillevirus branch. Although only one LCV genome, that of a Marseille-like virus, appears to be complete and on a single contig, several other genomes seem to be near complete, and overall, the LCV genomic data are sufficient to dramatically expand the pangenome of the PIM group, to add substantially to the NCLDV pangenome as well, and to reveal notable evolutionary trends. One of such trends is the apparent independent origin of giant viruses in more than one clade within both the Pithovirus and the Marseillevirus branches. Although this observation should be interpreted with caution, given the lack of fully assembled LCV genomes, it supports and extends the previous conclusions on the dynamic nature of NCLDV evolution (“genomic accordion”) that led to the independent, convergent evolution of viral gigantism in several, perhaps, even all NCLDV families (45, 48, 122, 123). Conversely, these findings are incompatible with the concept of reductive evolution of NCLDV from giant viruses as the principal evolutionary mode. Another notable evolutionary trend emerging from the LCV genome comparison is the apparent extensive gene exchange between Pitho-like and Marseille-like viruses, and members of the *Mimiviridae*. Finally, it is important to note that the LCV analysis reaffirms, on a greatly expanded dataset, the previously proposed monophyly of the PIM group of the NCLDV, demonstrating robustness of the evolutionary analysis of conserved NCLDV genes (28, 45). Furthermore, a congruent tree topology was obtained by gene content analysis, indicating that, despite the open pangenomes and the dominance of unique genes, evolution of the genetic core of the NCLDV appears to track the sequence divergence of the universal marker genes.

Like other giant viruses, several LCV encode multiple translation system components. Although none of them rivals the near complete translation systems encoded by Klosneuviruses (46), Orpheovirus (19), and especially, Tupanviruses (98), some are comparable, in this regard, to the Mimiviruses (45). The diverse origins of the translation system components in LCV suggested by phylogenetic analysis are compatible with the previous conclusions on the piecemeal capture of these genes by giant viruses as opposed to inheritance from a common ancestor (43, 45).

The 23 NCLDV genome bins reconstructed in the present study only represent a small fraction of the full NCLDV diversity as determined by DNA polymerase sequences present in marine sediments (Figure 1). Notably, sequences closely matching the sequences in the NCLDV genome bins were identified only in the Loki’s Castle metagenomes, not in TARA oceans water column metagenomes or Earth Virome sequences. Thus, the deep sea sediments represent a unique and unexplored habitat for NCLDVs. Further studies targeting deep sea sediments will bring new insights into the diversity and genomic potential of these viruses.

Identification of the host range is one of the most difficult problems in metaviromics and also in the study of giant viruses, even by traditional methods. Most of the giant viruses have been isolated by co-cultivation with model amoeba species, and the natural hosts remains unknown. Notable exceptions are the giant viruses isolated from marine flagellates *Cafeteria roenbergensis* (12) and *Bodo saltans* (35). The principal approach for inferring the virus host range from metagenomics data is the analysis of co-occurrence of virus sequences with those of potential hosts (124, 125). However, virtually no 18S rRNA gene sequences of eukaryotic origin were detected in the Loki’s Castle sediment samples, in a sharp contrast to the rich prokaryotic microbiota (61, 62). The absence of potential eukaryotic hosts of the LCV strongly suggests that these viruses do not reproduce in the sediments but rather could originate from virus particles that precipitate from different parts of the water column. So far, however, closely related sequences have not been found in water column metagenomes (Figure 1). The eukaryotic hosts might have inhabited the shallower sediments, and although they have decomposed over time, the resilient virus particles remain as a “fossil record”. Clearly, the hosts of these viruses remain to be identified. An obvious and important limitation of this work – and any metagenomic study – is that the viruses discovered here (we are now in a position to call the viruses without quotes, given the recent decisions of the ICTV) have not been grown in a host culture. Accordingly, our understanding of their biology is limited to the inferences made from the genomic sequence which, per force, cannot yield the complete picture. In the case of the NCLDV, these limitations are exacerbated by the fact that their genomic DNA is not infectious, and therefore, even the availability of the complete genome does not provide for growing the virus. The metagenomic analyses must complement rather than replace traditional virology and newer culturomic approaches.

Although the sediment samples used in this study have not been dated directly, determinations of sedimentation rates in nearby areas show that these rates vary between 1-5 cm per 1000 years (126, 127). With the fastest sedimentation rate considered, the sediments could be over 20,600 years old at the shallowest depth (103 cm). Considering that *Pithovirus sibericum* and *Mollivirus sibericum* were revived from 30,000 year old permafrost (17, 20), it might be possible to resuscitate some of the LCVs using similar methods. Isolation experiments with giant viruses from deep sea sediments, now that we are aware of their presence, would be the natural next step to learn more about their biology.

Regardless, the discovery of the LCV substantially expands the known ocean megavirome and demonstrates the previously unsuspected high prevalence of Pitho-like viruses. Given that all this diversity comes from a single site on the ocean floor, it appears clear that the megavirome is large and diverse, and metagenomics analysis of NCLDV from other sites will bring many surprises.

## Acknowledgements

We acknowledge the help from chief scientist R. B. Pedersen, the scientific party and the entire crew on board the Norwegian research vessel G.O. Sars during the summer 2008, 2010 and 2014 expeditions. We thank the Uppsala Multidisciplinary Center for Advanced Computational Science (UPPMAX) at Uppsala University and the Swedish National Infrastructure for Computing (SNIC) at the PDC Center for High-Performance Computing for providing computational resources. This work was supported by grants of the European Research Council (ERC Starting grant 310039-PUZZLE_CELL), the Swedish Foundation for Strategic Research (SSF-FFL5) and the Swedish Research Council (VR grant 2015-04959) to T.J.G.E. N.Y., YIW, and E.V.K. are funded through the Intramural Research program of the National Institutes of Health of the USA.

## Additional files

Additional File 1 – Supplementary binning methods and figures

Additional File 2 – LCV and LC virophage protein annotation

Additional File 3 – DNAp, MCP, A18hel, and translation protein trees

Additional File 4. – virophage genome maps

Additional File 5. – taxonomic breakdown of psi-BLAST hits retrieved with profiles created from selected PIM clusters (clusters of four or more proteins, less conserved NCLDV genes).

Additional File 6. – Repeats plots

Additional File 7. – Conserved promoter-like motif frequencies in selected LCV bins

Additional File 8. – Conserved promoter-like motifs in the LCMiAC01 and LCMiAC02 bins, and LCV virophages

Additonal File 9. – Conserved promoter-like motifs in pitho-like LCV.

Additonal File 10. – Conserved promoter-like motifs in Marseille-like and irido-like LCV.

More supplementary material: ftp://ftp.ncbi.nih.gov/pub/yutinn/Loki_Castle_NCLDV_2018/

